# Fitness and functional landscapes of the *E. coli* RNase III gene *rnc*

**DOI:** 10.1101/2022.11.01.514689

**Authors:** Ryan Weeks, Marc Ostermeier

## Abstract

How protein properties such as protein activity and protein essentiality affect the distribution of fitness effects (DFE) of mutations are important questions in protein evolution. Deep mutational scanning studies typically measure the effects of a comprehensive set of mutations on either protein activity or fitness. Our understanding of the underpinnings of the DFE would be enhanced by a comprehensive study of both for the same gene. Here, we compared the fitness effects and in vivo protein activity effects of ∼4,500 missense mutations in the *E. coli rnc* gene. This gene encodes RNase III, a global regulator enzyme that cleaves diverse RNA substrates including precursor ribosomal RNA and various mRNAs including its own 5’ untranslated region (5’UTR). We find that RNase III’s ability to cleave dsRNA is the most important determinant of the fitness effects of *rnc* mutations. The DFE of RNase III was bimodal, with mutations centered around neutral and deleterious effects, consistent with previously reported DFE’s of enzymes with a singular physiological role. Fitness was buffered to small effects on RNase III activity. The enzyme’s RNase III domain (RIIID), which contains the RNase III signature motif and all active site residues, was more sensitive to mutation than its dsRNA binding domain (dsRBD), which is responsible for recognition and binding to dsRNA. Differential effects on fitness and functional scores for mutations at highly conserved residues G97, G99, and F188 suggest that these positions may be important for RNase III cleavage specificity.

## Introduction

In the last ten years, comprehensive experiments on the effects of all single-codon missense mutations, called deep mutational scanning (DMS) (Fowler and Fields 2014), have been performed on a variety of proteins (Roscoe, et al. 2013; Firnberg, et al. 2014; Stiffler, et al. 2015; Mavor, et al. 2016; Wrenbeck, et al. 2017; Wu, et al. 2022). Many of these studies report on the mutational effects on protein properties. These studies often use non-native reporter assays, sometimes with the protein expressed in a non-native organism, to determine, for example, mutational effects on the affinity of an inhibitor against H1N1 influenza hemagglutinin (Whitehead, et al. 2012). These DMS studies show how changes to the amino acid sequence affect protein activity. Although these studies often are described as producing “fitness landscapes”, these landscapes are not fitness landscapes in the traditional sense of the word “fitness” (i.e. the ability of an organism to survive and procreate). In contrast, traditional fitness landscapes describe the effect of mutations on the fitness of the organism (Lind, et al. 2010; Roscoe, et al. 2013; Firnberg, et al. 2014; Lind, et al. 2016; Mavor, et al. 2016; Lundin, et al. 2017; Wrenbeck, et al. 2017; Hartman and Tullman-Ercek 2019; Asgari, et al. 2020) and are thus more relevant for protein evolution.

The distribution of fitness effects of mutations (DFE) governs a gene’s potential evolutionary paths. DFEs of missense mutations have been reported for diverse proteins such as enzymes, binding proteins, transcription factors, and ribosomal proteins (Lind, et al. 2010; Hietpas, et al. 2011; Jiang, et al. 2013; Melamed, et al. 2013; Roscoe, et al. 2013; Firnberg, et al. 2014; Stiffler, et al. 2015; Lind, et al. 2016; Mavor, et al. 2016; Lundin, et al. 2017; Noda-García, et al. 2019; Mehlhoff, et al. 2020; Wu, et al. 2022), although some of these studies have examined just a small fraction of the possible missense mutations. Nearly all DFEs have been bimodal, with peaks centered on fully deleterious and neutral effects. Notable exceptions include non-comprehensive mutational studies of two *S. typhimurium* ribosomal proteins (Lind, et al. 2010). The fractions of mutations observed in each peak of the DFEs is a measure of the protein’s mutational robustness, which has implications for a protein’s evolution. For many proteins, most mutations are neutral (Roscoe, et al. 2013; Firnberg, et al. 2014; Stiffler, et al. 2015; Lind, et al. 2016; Mavor, et al. 2016; Lundin, et al. 2017), but for others most are deleterious (Lind, et al. 2016). Important factors causing such differences in DFEs include the level of gene expression and the environment in which fitness is measured (Stiffler, et al. 2015; Adams, et al. 2016; Mavor, et al. 2016; Lundin, et al. 2017; Noda-García, et al. 2019). When expression is sufficiently high, fitness effects can be buffered to mutational effects that do not decrease activity below an activity threshold, as observed with *S. enterica* HisA and *S. cerevisiae* Hsp90 (Hietpas, et al. 2011; Jiang, et al. 2013; Lundin, et al. 2017).

Recent studies of *S. cerevisiae URA3* (Wu, et al. 2022) and *E. coli TEM-1* β-lactamase (Stiffler, et al. 2015; Mehlhoff, et al. 2020) indicate that a gene’s essentiality in the growth environment also shapes the DFE, with mutations less deleterious to protein function being tolerated in environments where the gene is less essential for growth. Other properties, such as the role of the protein in the cell and the centrality of the protein to the cell’s function may affect the DFE. The fitness effects of a small sampling of mutations in *S. typhimurium* ribosomal protein genes, genes whose loss causes a 75-80% reduction in fitness, suggested that important genes with central roles in the cell may be quite robust to mutation (Lind, et al. 2010). Conceivably, the type of protein function may also play a role. A recent study of T4 bacteriophage sliding clamp and its clamp loader found that the enzymatic domain of the clamp loader was the least tolerant to mutation with a bimodal DFE, whereas the sliding clamp, which has a structural role, was robust to mutation (Subramanian, et al. 2021). The tolerance of the collar domain of the clamp loader, which is part of the enzyme but lacks catalytic activity on its own, was intermediate between that of the enzyme domain and the sliding clamp.

Many DFE studies examine only a limited subset of mutations (e.g. randomly-generated mutations or those made to a certain region of the protein important to function) and thus do not provide a comprehensive picture of the DFE. To date, only a few comprehensive fitness landscapes have been reported in native organisms including *TEM-1* β-lactamase (Firnberg, et al. 2014; Stiffler, et al. 2015; Mehlhoff, et al. 2020), *folA* encoding dihydrofolate reductase (DHFR)(Thompson, et al. 2020), *URA3* encoding an orotidine 5-phosphate decarboxylase (Wu, et al. 2022), and ubiquitin (Roscoe, et al. 2013; Mavor, et al. 2016). *TEM-*1 and *URA3* encode enzymes that have a singular function in the cell (e.g. TEM-1 degrades an antibiotic); conversely, ubiquitin, a small binding protein with no catalytic activity, has a wide array of regulatory roles in the cell. Do proteins with multiple cellular roles have a different level of mutational tolerance than proteins with singular roles? One could imagine that such proteins would be more sensitive to mutations if their multiple cellular roles add additional constraints on protein sequence. Alternatively, they could be less sensitive to mutations because their multiple roles caused them to evolve to be more robust. The fitness landscape for ubiquitin (Roscoe, et al. 2013; Mavor, et al. 2016) suggests that essential proteins involved in multiple processes might be more tolerant to mutations, while the fitness landscape of all single-codon mutations of one domain of the *S. cerevisiae* RNA poly(A)-binding protein Pab1 (Melamed, et al. 2013) suggests the opposite may be true. Alternatively, ubiquitin’s tolerance to mutation may be due to its high stability (Makhatadze, et al. 1998).

An important question in evolutionary biology is what causes the fitness effects of mutations. While it is expected that fitness will depend on the protein’s properties and activity, the functional form of this dependence is expected to be nonlinear (in part because of buffering), has only been studied for a limited number of mutations (Bershtein, et al. 2012; Rodrigues, et al. 2016; Sarkisyan, et al. 2016; Thompson, et al. 2020), and may well take different forms for different proteins. In addition, the collateral fitness effects of mutations (i.e. those effects that do not originate from changes in the ability of the gene to perform its physiological function) may play an important role (Mehlhoff, et al. 2020). A variety of DMS studies on the functional effects of mutations indicated that their distribution is also bimodal (Bandaru, et al. 2017; Heredia, et al. 2018; Shah and Kuriyan 2019; Subramanian, et al. 2021). To our knowledge, there has been no study to measure the effects of all mutations on both fitness and protein activity for the same gene. Such a study would add to our overall understanding of how fitness depends on protein function.

Here, we chose to study the fitness and protein activity landscapes of *E. coli* ribonuclease III (RNase III), a dsRNA-specific, Mg^2+^ dependent endonuclease that is involved in many processes in the cell including the maturing of rRNA and post-transcriptionally regulating numerous genes including its own (Dunn and Studier 1973; Gitelman and Apirion 1980; Sim, et al. 2010; Stead, et al. 2011; Court, et al. 2013; Sim, et al. 2014; Gordon, et al. 2017; Altuvia, et al. 2018). Members of the RNase III family are found in organisms from bacteria to humans and are characterized by the presence of a ten-residue signature motif and by their ability to cleave dsRNA. Although the functional roles of RNase III family members differ, sequence conservation of the RIIIDs and dsRBDs remains high. Within eukaryotes, including humans, Drosha and Dicer are complex RNase III enzymes that are responsible for processing small RNAs involved with RNA interference (Filippov, et al. 2000; Wu, et al. 2000; Bernstein, et al. 2001; Lee, et al. 2003).

Within the RNase III enzyme family, *E. coli* RNase III is one of the simplest enzymes. Despite its involvement in many processes, *E. coli* RNase III is not essential (Apirion and Watson 1974; Gegenheimer and Apirion 1975; Babitzke, et al. 1993; Hauk, et al. 2022), but cells lacking RNase III grow about 20-27% slower (Babitzke, et al. 1993). Monomers of RNase III homodimerize to create two identical active sites surrounding target dsRNA and catalyzes the two strand-cleavage reactions (Blaszczyk, et al. 2001; Gan, et al. 2006; Court, et al. 2013; Altuvia, et al. 2018). Each RNase III monomer is composed of an N-terminal RNase III domain (RIIID), which contains the RNase III signature motif and all active site residues, and a C-terminal dsRNA binding domain (dsRBD) which is responsible for recognition and binding to dsRNA (Court, et al. 2013). Acting on its most abundant substrate, RNase III regulates ribosome formation through maturation of rRNA transcribed from rRNA operons (Kaczanowska and Rydén-Aulin 2007; Deutscher 2009; Court, et al. 2013). RNase III also activates and represses gene expression through dsRNA cleavage of an array of different mRNA messages (Sim, et al. 2010; Stead, et al. 2011; Adkar, et al. 2012; Gordon, et al. 2017; Altuvia, et al. 2018). For activation, RNase III cleavage can remove barriers that block translation, such as freeing the ribosome binding site in *adhE* mRNA for complex with the ribosome (Aristarkhov, et al. 1996).

Conversely, RNase III cleavage can make mRNA more susceptible to degradation, as is exemplified by the negative feedback that regulates RNase III activity in cells. When cellular RNase III levels are high, cleavage of its own 5’ untranslated region (5’-UTR) leaves the *rnc* mRNA vulnerable to RNase E degradation thus downregulating RNase III levels (Bardwell, et al. 1989; Régnier and Grunberg-Manago 1989, 1990; Matsunaga, Dyer, et al. 1996; Matsunaga, Simons, et al. 1996; Nicholson 1999). In addition to its relevance for protein evolution, a comprehensive fitness and protein activity landscape of RNase III would allow for a more complete picture of how the sequence of *E. coli* RNase III is related to its function, as no crystal structure exists of the *E. coli* enzyme.

## Results

### Cells lacking a functional RNase III grow 25% slower

We previously demonstrated that when RNase III is placed under the pBAD promoter on a plasmid, RNase III is expressed even in the absence of the promoter’s inducer, arabinose (Hauk, et al. 2022). Under these leaky expression conditions, the cleavage pattern of the *rnc* 5’UTR (a native substrate) is similar to that when RNase III is chromosomally-expressed (Hauk, et al. 2022). Thus, we presume the RNase III expression levels were approximately physiological. In those experiments, the plasmid-encoded RNase III lacked it’s native 5’UTR. In our experiments here, we integrated the native *rnc* 5’-UTR in place of the pBAD 5’-UTR on pRnc-gGFP (Hauk, et al. 2022) to make pRnc2-gGFP (**fig. 1A**) to better reflect the native autoregulation of RNase III.

**Figure 1:**
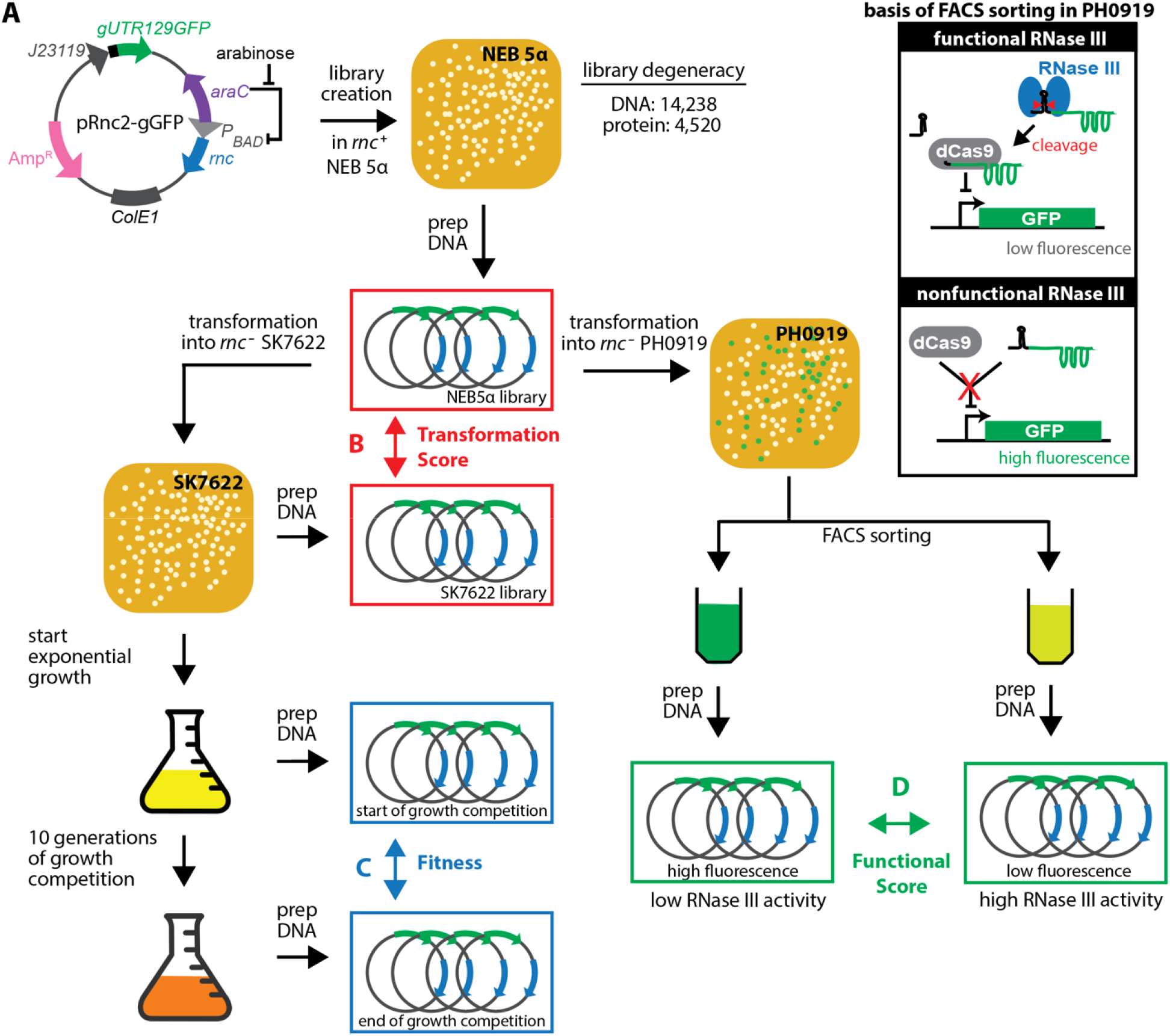
Experimental design for measuring the fitness, transformation, and functional landscapes of RNase III. (**A**) A comprehensive site-saturation library of the coding region of the RNase III gene was created on plasmid pRnc2-gGFP in *rnc*^*+*^ NEB5*α* cells. (**B**) Transformation scores were determined from the change in mutation frequency in the library before and after transformation into *rnc*^*–*^ SK7622 cells. (**C**) Fitness values were determined from the change in mutation frequency in SK7622 cells after ten generati ns of exponential growth in LB media. To measure (**D**) functional scores, the library was transformed into *rnc*^*–*^ PH0919 cells that contain a fluorescent reporter for in vivo RNase III cleavage activity (Hauk, et al. 2022). These cells were sorted by FACS into high and low fluorescent populations. The functional score was determined by the difference in mutation frequency between the two populations relative to wild type.

We first confirmed that expression of RNase III in this manner restored wildtype growth rates in cells lacking a functional *rnc* gene on the chromosome. We expressed wildtype RNase III, the nuclease-null RNase III E117K, and the empty vector (EV) in MG1693 and SK7622 (MG1693 *rnc* ^−^) cells. RNase III E117K lacks the ability to catalytically cleave dsRNA but maintains the ability to specifically bind with dsRNA (Dasgupta, et al. 1998; Sun and Nicholson 2001). We measured the growth rate at 37°C of exponentially growing cells in liquid lysogeny broth (LB) media containing 0.2% glucose (a repressor of the pBAD promoter) and ampicillin (Amp) for plasmid maintenance (**supplementary fig. S1**, Supplementary Material online). The SK7622 cells expressing RNase III-E117K and EV grew approximately 25% slower than cells with chromosomally-expressed RNase III (*P*<0.0001, 0.0003 by unpaired Student’s *T*-test for E117K and EV, respectfully). However, no significant difference in growth rate was observed between SK7622 cells expressing RNase III and MG1693 cells expressing any of the three constructs. As inducing higher expression levels of RNase III by adding typical amounts of arabinose results in irregular sized bacteria with a diminished growth rate (Hauk, et al. 2022), we interpret our no arabinose growth conditions as producing approximately physiological expression levels of Rnase III. This is also consistent with the observation that *E. coli* growth rate is not affected by 0.1 to 10-times the endogenous levels of RNase III (Sim, et al. 2010). However, we did not quantify expression by measuring RNase III mRNA or protein levels as our experimental set up was already non-native by virtue of *rnc* being on a multi-copy plasmid instead of the chromosome.

### Fitness landscape measured in growth competition experiments

We constructed a library of all single-codon substitutions in *rnc* on the pRnc2-gGFP plasmid in the *rnc*^+^ cloning strain NEB5α. We used NEB5α as a library construction strain to avoid biasing the library by the fitness effects of RNase III mutations during library construction. We purified the library plasmid DNA, and deep sequencing revealed it consisted of nearly all single-codon amino acid mutations except for poor coverage at one position: I6. This plasmid library was transformed into SK7622 cells, which lack a functional RNase III gene. In two replica experiments, exponentially growing SK7622 cells containing the library were subject to a growth competition experiment in LB media for 10 generations (**fig. 1C**). We determined the frequency of each *rnc* mutant by deep-sequencing at the beginning and end of the competition. From these frequencies, we determined relative fitness (*w*) values, corresponding to the mean growth rate of cells containing a mutant RNase III relative to the mean growth rate of cells containing alleles synonymous to wildtype (Mehlhoff and Ostermeier 2020). We utilize the frequency of wildtype synonymous alleles instead of the frequency of wildtype because wildtype synonyms occurred more frequently in the library and wildtype sequencing counts are more prone to being affected by the artifact of PCR template jumping (Pääbo, et al. 1990; Kebschull and Zador 2015) during the preparation of barcoded amplicons for deep-sequencing.

Curiously, although fitness values correlated between the two replica experiments (**supplementary fig. S2**, Supplementary Material online), the mean fitness in replica 1 (0.952) was higher than that of replica 2 (0.923) and over twice as many mutants in replica two were deemed significantly deleterious at *P*<0.01 (48% vs 21%). A comparison of the two DFEs shows that a subset of mutants that were near neutral in replica 1 became slightly deleterious in replica 2 and those mutations deleterious in replica 1 became, on average, slightly more deleterious in replica 2 (**supplementary fig. S2**, Supplementary Material online). For unknown reasons, it appears that the RNase III activity required for wildtype growth rates was more stringent during replica 2. In replica 2 we observed that a wildtype reference monoculture grown at the same time as the libraries grew 3% faster and the libraries grew 1.1-1.7% faster than in replica 1 (**supplementary fig. S3**, Supplementary Material online), indicating that something was different about the growth conditions between the two replica experiments, perhaps the temperature was slightly higher in replica 2. We chose to present our fitness values as the weighted mean of the two replicas.

We obtained fitness data for 99.58% (4501/4520) of single-codon mutation in RNase III (**fig. 2A**). A total of 44.8% (2006/4479) of mutations nominally had fitness values under 0.98, with 54.6% being near-neutral (fitness values between 0.98 and 1.02). A total of 19.0% (*P* <0.01, *Z*-test) and 13.6% (*P*<0.001) of the mutations were deleterious in both replica experiments. As expected, regions of predicted secondary structure, namely the α-helices in the RIID and dsRBD domains, were more prone to deleterious mutations than loop regions between secondary structural elements. Mutations to the most exchangeable amino acid (Yampolsky and Stoltzfus 2005) tended to have smaller fitness effects with a median fitness of 0.994 ± 0.054 (**supplementary fig. S4**, Supplementary Material online).

**Figure 2:**
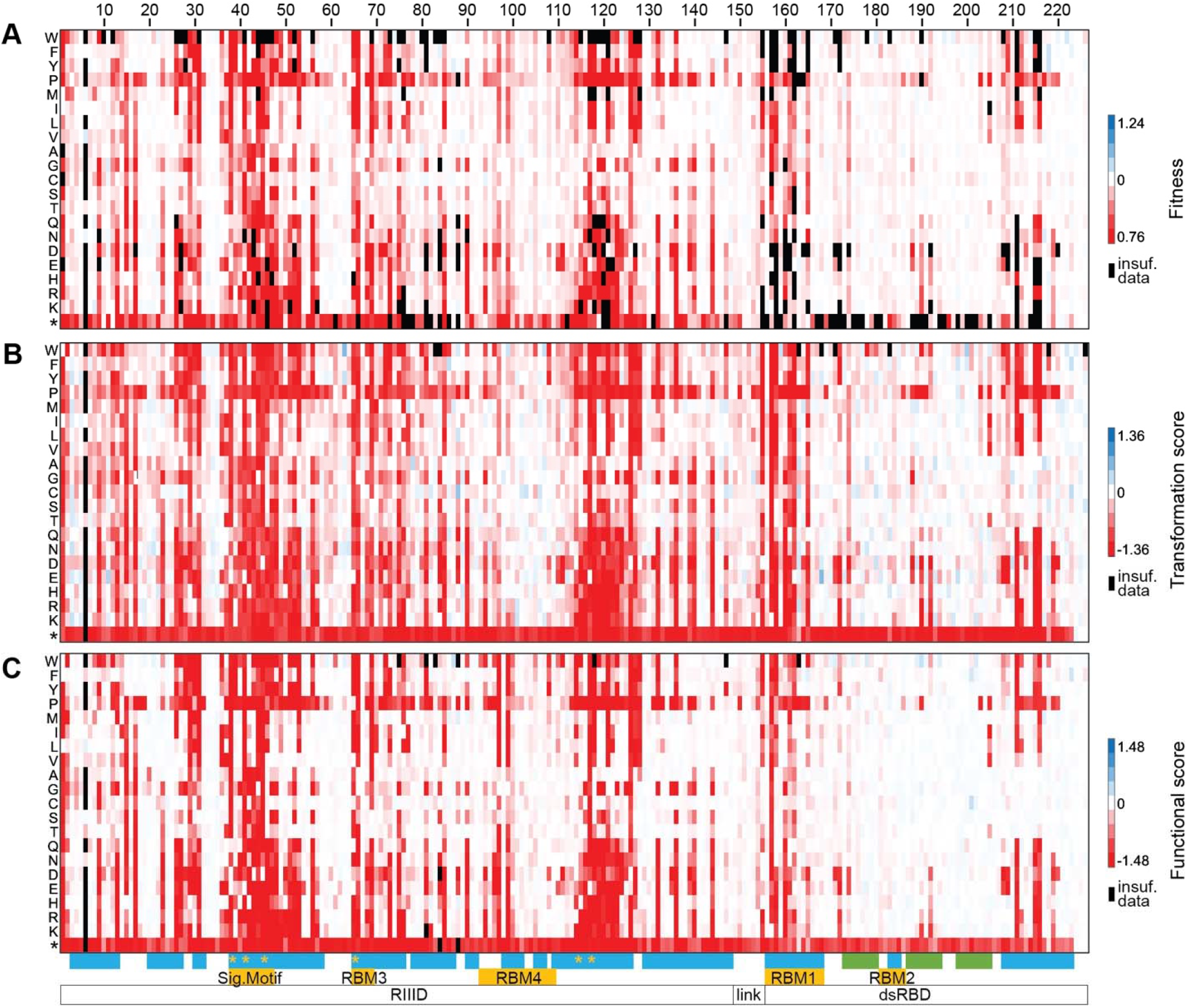
Fitness, transformation scores, and functional scores are qualitatively alike. Heat maps show the (**A**) Fitness, (**B**) transformation score, and (**C**) functional score landscapes of *E. coli* RNase III. Red is deleterious, white is neutral, blue is beneficial, and black indicates insufficient data due to low sequencing counts. The shading for each landscape was set such that full red represented the average value corresponding to complete loss of RNase III activity. The amino acid substitutions are shown on the left (* = nonsense mutation). Fitness and functional scores are reported as the weighted mean of two replica experiments. Transformation score is a single experiment. At the bottom are secondary structure elements (*α*-helices in blue and *β*-strands in green); functional motifs in orange including the signature motif, the four RNA binding motifs (RBM), and the six negatively charged active site residues (orange *); and the RIIID and dsRBD domains joined by the linker. The values and errors of all fitness, transformation scores, and functional scores can be found in **supplementary data S1-S3**, Supplementary Material online.

Surprisingly, the distribution of fitness effects (DFE) showed an apparent unimodal distribution centered at neutral effects and a long tail of deleterious mutations (**fig. 3A**). Typically, DFE’s are bimodal (Lind, et al. 2010; Peris, et al. 2010; Roscoe, et al. 2013; Firnberg, et al. 2014; Lind, et al. 2016; Lundin, et al. 2017; Mehlhoff and Ostermeier 2020), with one peak centered at neutral effects and the other corresponding to fitness values for an inactive protein (Lind, et al. 2010). Similar unimodal distributions have been observed in ribosomal proteins shown to be highly robust to mutations (Lind, et al. 2010; Lind, et al. 2016). However, the tail of deleterious mutations was very broad, and we noticed that mutations with low fitness measurements tended to have lower than average sequencing counts at the start of the growth competition. These low counts resulted in a decreased ability to precisely measure their fitness effect as reflected in the uncertainty and *p*-values, which are inversely proportional to the sequencing counts. Furthermore, many nonsense mutations had fitness values well-above or well-below the expected value 0.75, the fitness value of cells lacking functional RNase III. Instead, nonsense mutations had a wide range of fitness values and a median fitness of 0.839 ± 0.109 (**supplementary fig. S4**, Supplementary Material online). Fitness values of nonsense mutations typically had large uncertainties stemming from low counts at the start of the experiment. We hypothesized that cells with deleterious RNase III mutations might not survive the transformation step as frequently, might form smaller colonies on the transformation plate, and would grow slower or have a longer lag phase in the pre-exponential incubation period prior to the growth competition phase of the experiment. This imprecision in the measurement of deleterious effects would broaden the expected peak corresponding to inactivating mutations, leading to a false appearance of a unimodal distribution. We sought another method to more accurately measure the fitness effects of mutations of deleterious mutations.

**Figure 3:**
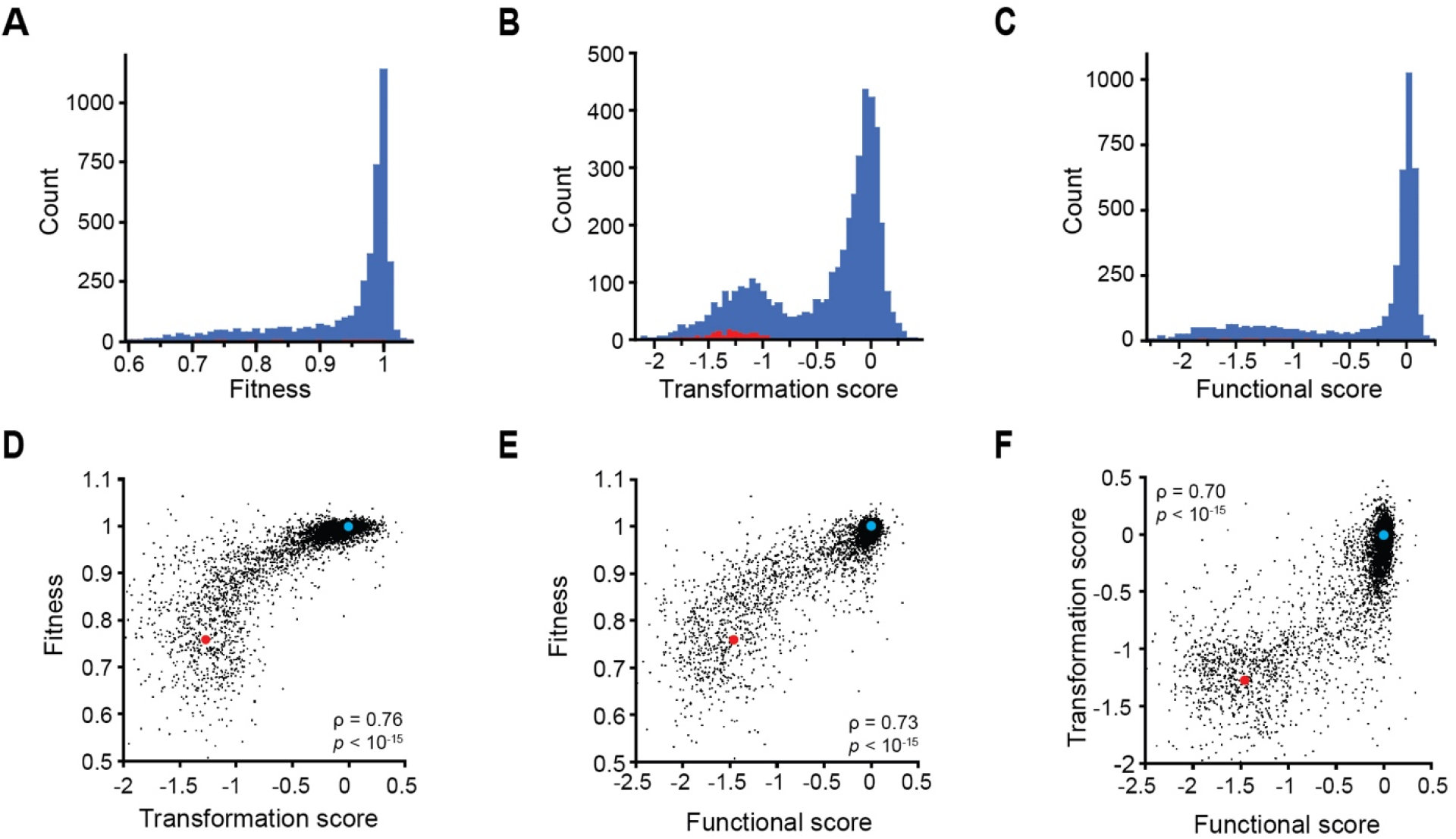
The distribution of mutational effects for transformation and functional scores are bimodal and the three fitness metrics correlate with each other. Histograms of (**A**) the weighted mean fitness, (**B**) the transformation score, and (**C**) the weighted mean functional score for missense (blue) and nonsense (red) mutations. The distribution of nonsense mutations is too broad to observe in A and C. The distribution of nonsense mutation functional scores can be seen in **supplementary fig. S6B,C**, Supplementary Material online. (**D**-**F**) Relationship between fitness, transformation score, and functional score. The values for wildtype (blue) and nonfunctional RNase III (red) are shown for reference. The value for nonfunctional RNase III fitness is the growth rate of cells with the M1S mutation (i.e. start codon replaced with TCT) relative to that of wildtype. The values for nonfunctional RNase III transformation and functional scores are the weighted mean of all mutants with a non-functional start codon.

### Deleterious fitness effects observed through mutant-dependent differences in transformation efficiency

We deep sequenced the NEB5α and SK7622 libraries to assess whether the transformation step negatively affected the frequency of deleterious mutants in the library (**fig. 1B**). From the frequencies of each mutant in each library, we determined transformation scores (*w*_*t*_) corresponding to the log of the ratio of the enrichment of cells containing a mutant relative to the enrichment of cells containing alleles synonymous for the wildtype (**fig. 2B**). Transformation fitness scores are presented in log form, such that deleterious mutants with a score of -1 are ten-fold depleted in the SK7622 library relative to wildtype synonyms.

The results indicated that transformation of the library into SK7622 altered the frequencies of certain types of mutations. If transformation effectiveness was unaffected by RNase III function, one would expect that transformation fitness scores would show a random distribution centered about a value of zero regardless of RNase III function. However, this was not the case as nonsense mutations (excluding the last three codons) had a weighted mean transformation score of -1.36, but wildtype synonyms had a weighted mean transformation fitness score of 0.00 (**supplementary fig. S5**, Supplementary Material online). Nonsense mutation frequency diminished from 4.0% in the NEB5α library (close to the expected frequency of 4.7% (3/64)) to nearly tenfold less, representing only 0.55% of mutations in the SK7622 library. Conversely, residues mutated to those that are considered the most exchangeable were enriched from 4.3% in in our NEB5α library to 5.6% in the SK7622 library.

The distribution of transformation fitness scores was bimodal, with a neutral peak near 0 and a deleterious peak near -1.25 (**fig. 3B**). The landscapes of fitness and transformations scores looked qualitatively similar (**fig. 2A,B**), and fitness correlated with transformation scores (*ρ* = 0.76, *p* < 10^−15^, **fig. 3D**). Thus, transformation scores likely report on many of the same fitness-impacting mutational effects that occur when the cells grow exponentially in liquid media. Transformation scores represent the relative ability of *rnc* ^−^ cells to acquire the plasmid-encoded mutant RNase III allele and grow on solid media. We conclude that the apparent absence of an obvious peak for loss of function mutations in the RNase III fitness landscape is just an artifact of the inability of the fitness measurements to precisely measure deleterious fitness effects.

Interestingly, a sizable subset of mutants had deleterious transformation scores, but near neutral fitness values for exponential growth (**fig. 3D**). This difference may have several, non-mutually exclusive reasons. First, this difference may arise because certain functions of RNase III become more important for fitness during the stress of transformation or for growth on solid media. Second, the stress of transformation may make the cells more sensitive to small changes in RNase III catalytic activity, changes that are too small to have fitness consequences during exponential growth in liquid media.

Finally, transformation scores might be more sensitive to small deleterious effects of mutations on RNase III activity because transcription of the RNase III gene starts after transformation of the plasmid into the cell. Thus, there would be a lag before the RNase III protein reaches steady state levels and can ameliorate the negative fitness effects experienced by the *rnc*^*–*^ cells prior to transformation. Regardless of the mechanism(s), transformation scores are an important added dimension for understanding the fitness consequences of RNase III mutations in addition to being a more precise measure of the fitness effects of deleterious mutants.

### Functional landscape for RNase III cleavage of a native substrate in vivo

How fitness depends on protein properties is an important question for the study of fitness landscapes. One difficulty in addressing this question is the lack of methods for high-throughput measurements of protein activity in vivo. We developed a high throughput system that reports the ability of RNase III to cleave a native RNase III substrate: the 5’ UTR of its own mRNA message (Hauk, et al. 2022) (upper right inset of **fig. 1**). This system utilizes an *E. coli* strain in which chromosomally-integrated GFP is transcriptionally-repressed by nuclease-null dCas9 when a GFP-targeting guide RNA (gGFP) is co-expressed (Qi, et al. 2013). Instead of using gGFP, we engineered a hybrid gRNA in which a fragment of the *rnc* 5’-UTR (nucleotides 1-129) is fused to the 5’ end of gGFP (gUTR129GFP) (Hauk, et al. 2022). In the absence of RNase III, the gUTR129GFP is compromised in its ability to function with dCas9 to repress GFP expression, presumably by sterically blocking complexation with dCas9. However, RNase III cleavage of the gUTR129GFP causes release of the gGFP, which results in dCas9-mediated GFP repression. We showed that this fluorescent-reporter system allowed us to sort cells using flow cytometry based on RNase III activity (Hauk, et al. 2022). Cells lacking RNase III activity could be enriched from an excess of cells containing RNase III activity and vice versa.

Here, we use this system to measure a functional score for all mutations. We transformed the NEB5α library into the fluorescent-reporter, *rnc*^*–*^ strain PH0919, which co-expresses dCas9 from a separate, compatible plasmid. The library was sorted into two populations by FACS: cells with higher fluorescence and cells with lower fluorescence (**fig. 1D**). The gates for selecting the two populations were determined based on the forward scatter versus fluorescence plots of cells expressing wildtype RNase III and cells expressing the E117K inactive mutant, as previously demonstrated (Hauk, et al. 2022). The frequency of each mutant in each population was determined by deep-sequencing. The enrichment of each mutant was defined as the ratio of the frequency of each mutant in the low-fluorescence, active population to its frequency in the high-fluorescence, inactive population. From these enrichments, we determined the functional scores (*w*_*f*_) corresponding to the log of the ratio of the enrichment of cells containing a mutant relative to the enrichment of cells containing alleles synonymous for the wildtype.

Functional scores were calculated in two replica experiments, but the range of values in the first replica was larger than in the second (**supplementary fig. S6A-C**, Supplementary Material online). For example, non-amber nonsense mutations within the RIIID had a weighted mean functional score of -1.85 in the first replica and -0.80 in the second. We believe the difference in ranges might have resulted from differences in the length of time between transformation and the sorting experiment, differences in the precision in placing the sorting gates, and differences in the efficiency of sorting. Although the two replicas had different ranges, the landscapes were qualitatively similar (**supplementary fig. S6A,D**, Supplementary Material online), so we present the functional scores as the weighted mean of the two replicas (**fig. 2C**). As observed for fitness values, deleterious mutants tended to have lower counts, which negatively impacted the precision of the functional scores of deleterious mutants. However, the problem was not as severe as in the fitness measurements because flow cytometry was performed on cells from the colonies on the transformation plates (i.e. no additional growth period occurred, unlike in the growth competition experiment).

The functional score represents how effective a mutant is at cleaving a native substrate, albeit in a non-native context. As observed with transformation fitness scores, the distribution for functional scores is bimodal, with peaks centered around -1.5 and 0 (**fig. 3C**). The bimodal nature of the distribution is best seen in each replica individually (**supplementary fig. S6B,C**, Supplementary Material online) as the averaging of the two replicas further broadened the lower peak beyond the broadening caused by the depletion of deleterious mutants during transformation. Functional scores positively correlated with fitness values (*ρ* = 0.73, *p* < 10^−15^, **fig. 3E**). Similarly, mutations with low functional scores tended to have low transformation scores (*ρ* = 0.70, *p* < 10^−15^, **fig. 3F**).

### Fitness is buffered to small deleterious effects on cellular RNase III activity

We constructed ten mutants of RNase III to accurately quantify their fitness and functional effects. The ten mutations include six to active site residues: E38A, E38V, E38Q, E65P, D114G, and D114R (Blaszczyk, et al. 2001; Sun, et al. 2004; Zhang, et al. 2004; Gan, et al. 2006; Xiao, et al. 2009). Other variants include D155E at the end of the flexible linker, expected to sterically interfere with RNase III binding to dsRNA (Inada and Nakamura 1995; Gan, et al. 2006), as well as F188D and C192D within β-strands in the dsRBD. We replaced the start codon with TCT, which cannot function as an alternative start (Hecht, et al. 2017), for a negative “no RNase III” control. We subjected these ten mutants to more in-depth experiments performed at the individual variant level (**Table 1**).

**Table 1:** Properties of individually studied mutants of *E. coli* RNase III.

| Variant | Fitness<sup>a,b</sup> | Transformation score<sup>c</sup> | Functional score |  |  | Fluor./OD<sup>e,b</sup> | Gel cleavage (%)<sup>b</sup> |
| --- | --- | --- | --- | --- | --- | --- | --- |
|  |  |  | Mean<sup>d</sup> | Replica 1<sup>c</sup> | Replica 2<sup>c</sup> |  |  |
| Wildtype | 1.00 ± 0.01 | 0.00 | 0.00 | 0.00 | 0.00 | 458 ± 78 | 69 ± 2 |
| E65P | 1.00 ± 0.01 | -0.26<br>+0.02/-0.02 | -0.09 | 0.04<br>+0.02/-0.02 | -0.13<br>+0.01/-0.01 | 401 ± 34 | 37 ± 7 |
| D114G | 0.99 ± 0.01 | -0.13<br>+0.03/-0.03 | -0.04 | 0.07<br>+0.03/-0.04 | -0.07<br>+0.01/-0.01 | 383 ± 5 | 48 ± 5 |
| E38Q | 0.99 ± 0.01 | -0.19<br>+0.02/-0.02 | -0.11 | -0.14<br>+0.02/-0.02 | -0.05<br>+0.01/-0.01 | 340 ± 6 | 47 ± 8 |
| D114R | 0.97 ± 0.01 | -0.57<br>+0.03/-0.03 | -0.02 | 0.01<br>+0.05/-0.05 | -0.03<br>+0.01/-0.01 | 349 ± 37 | 17 ± 4 |
| D155E | 0.90 ± 0.01 | -1.10<br>+0.08/-0.09 | -0.36 | -0.09<br>+0.04/-0.05 | -0.45<br>+0.05/-0.06 | 371 ± 22 | 13 ± 5 |
| E38A | 0.88 ± 0.01 | -1.05<br>+0.03/-0.03 | -1.47 | -1.51<br>+0.03/-0.03 | -0.46<br>+0.04/-0.04 | 543 ± 51 | 15 ± 4 |
| F188D | 0.83 ± 0.01 | -1.26<br>+0.08/-0.10 | -0.13 | -0.17<br>+0.07/-0.08 | 0.01<br>+0.04/-0.04 | 303 ± 8 | 27 ± 6 |
| E38V | 0.80 ± 0.01 | -1.03<br>+0.03/-0.03 | -1.55 | -1.58<br>+0.02/-0.02 | -0.65<br>+0.05/-0.06 | 747 ± 25 | 6 ± 4 |
| C192C | 0.80 ± 0.01 | -1.66<br>+0.18/-0.31 | -0.91 | -0.96<br>+0.06/-0.07 | -0.63<br>+0.16/-0.25 | 841 ± 145 | 13 ± 5 |
| TCT<sup>f</sup> | 0.76 ± 0.01 | -0.71<br>+0.07/-0.09 | -2.33 | -2.40<br>+0.15/-0.23 | -0.88<br>+0.20/-0.38 | 938 ± 54 | 7 ± 4 |
| ΔdsRBD<sup>g</sup> | 0.75 ± 0.01 | n.d.<sup>h</sup> | n.d. | n.d. | n.d. | 786 ± 95 | 2 ± 2 |
| E117K | 0.71 ± 0.01 | -1.28<br>+0.15/-0.23 | -2.18 | -2.20<br>+0.01/-0.01 | -0.84<br>+0.01/-0.02 | 1074 ± 36 | 3 ± 3 |

<sup>a</sup> Fitness determined as the growth rate relative to wildtype. Variants are ordered based on this value.
<sup>b</sup> Error is the standard deviation ( $n=3$ )
<sup>c</sup> Error is the standard deviation determined from the DMS sequencing counts
<sup>d</sup> Mean of replica 1 and replica 2 is weighted based on each replica's variance
<sup>e</sup> Fluorescence/OD-600nm of bacterial culture
<sup>f</sup> The M1 ATG start codon is replaced by the codon TCT, which does not function for translation initiation
<sup>g</sup> Q153\*/K154\*/D155\*
<sup>h</sup> n.d. = not determined

We measured the growth rates of SK7622 cells expressing each mutant under conditions identical to those in which we measured the fitness values by DMS. We calculated the relative fitness of these cells by dividing their growth rate by the growth rate of cells expressing the unmutated RNase III. A comparison of these fitness values to their transformation scores further illustrated that transformation score is a reasonable proxy for fitness in liquid media (**fig. 4A**).

**Figure 4:**
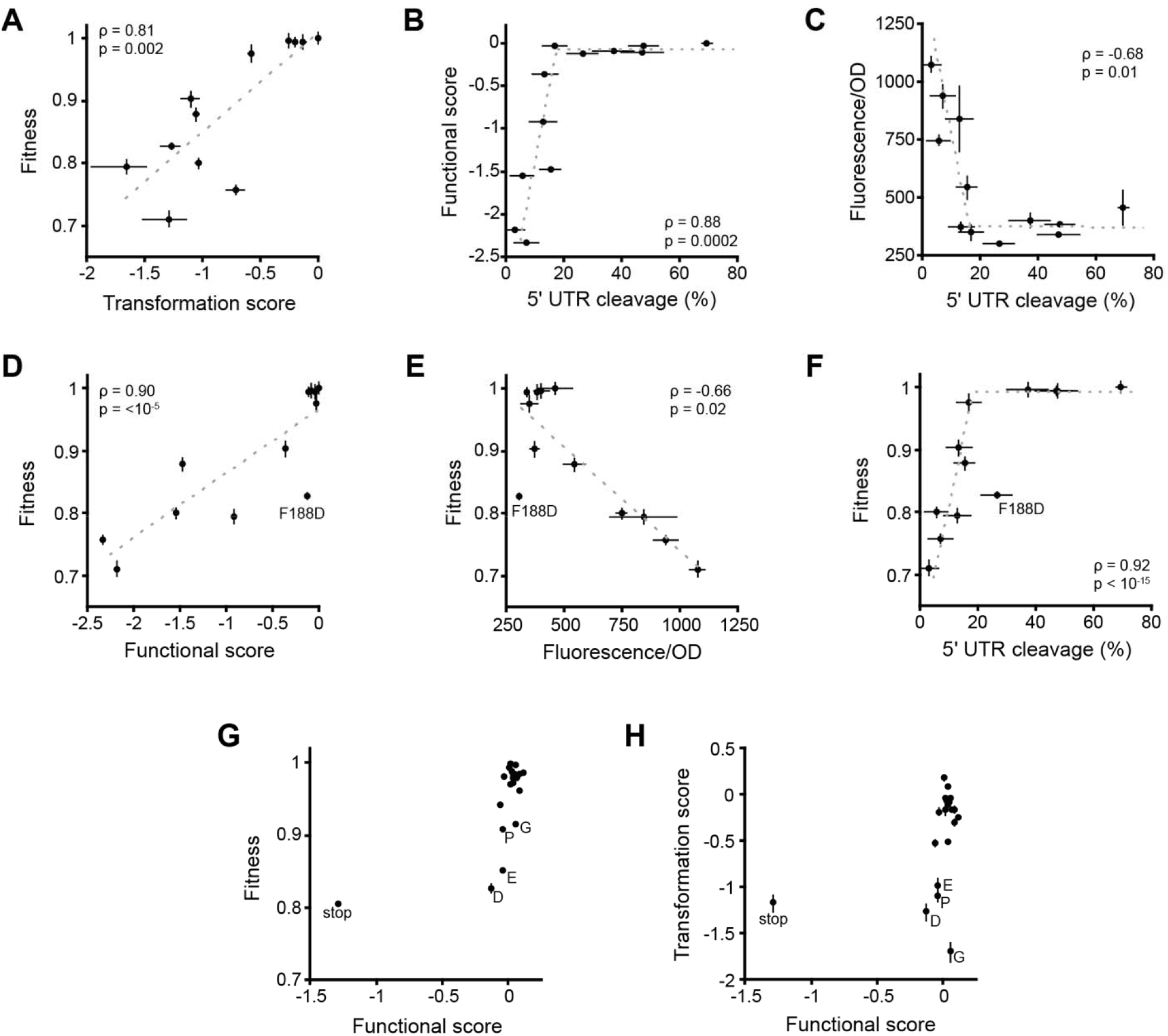
Relationship between function and fitness metrics for the select mutants of Table 1. (**A**) Fitness correlates with transformation score. (**B, C**) The functional score and the fluorescence activity assay are buffered against small changes in RNase III catalytic activity. (**D, E**) Fitness correlates with functional score and the fluorescence activity assay. (**F**) Fitness is buffered against small changes to RNase III catalytic activity. (**G, H**) Effects of all mutations at F188. Several mutations at F188 with deleterious effects on fitness and transformation score are identified. Fitness was determined by measuring the growth rate relative to wildtype (error bars are the standard deviation *n*=3) except for panel G where the fitness values for all mutants except F188D are the weighted mean values from the DMS experiment. Primer extension assays (**supplementary fig. S7**, Supplementary Material online) were used to determine the % cleavage of the 5’UTR stem loop of *rnc* in gUTR129GFP (error bars are the standard deviation *n*=3). The fluorescence/OD of bacterial cultures reports on RNase III cleavage of the 5’UTR stem loop of *rnc* in gUTR129GFP (error bars are the standard deviation *n*=3). Dashed lines are linear fits except for panels B, C, and F, in which case they are guides for the eye. Analogous graphs for each replica of the functional score (as opposed to the weighted mean used in panels B, D, G, and H) are provided as **supplementary fig. S8**, Supplementary Material online.

We also performed two RNase III functional assays on cultures of our exponentially-growing fluorescent reporter PH0919 cells expressing the mutants: 1) we measured the fluorescence of the cells, and 2) we used a primer extension assay to determine the gUTR129GFP cleavage efficiency in vivo (**supplementary fig. S7**, Supplementary Material online). A comparison of the cleavage assay to the functional score (**fig. 4B**) and to the fluorescent assay (**fig. 4C**) indicated that these fluorescence-based functional assays do not detect reductions in gUTR129GFP cleavage until it is reduced below a certain threshold. Below this threshold, functional scores decrease rapidly, and fluorescent values increase rapidly. Fitness was also buffered against deleterious effects on the ability of RNase III to cleave its own 5’UTR (**fig. 4F**). This buffering is expected from the known feedback control mechanism through which RNase III regulates its own expression by cleaving its own mRNA message (Bardwell, et al. 1989; Matsunaga, Dyer, et al. 1996; Matsunaga, Simons, et al. 1996). Such buffering could help ensure that the cell has an excess of RNase III activity above that necessary for wildtype growth rate. While we do not expect that our expression level is precisely physiological, we do observe sufficient expression to fully rescue the growth deficiency caused by loss of RNase III (**supplementary fig. S1**, Supplementary Material online) but do not have the excess expression that would decrease growth rate (Sim, et al. 2010) and cause abnormal cell morphology (Hauk, et al. 2022). However, if our expression level is much higher than the native level, excess buffering of fitness to small functional effects may result.

### Mutations at F188 differentially effect fitness and cleavage of the *rnc* 5’UTR

RNase III has many substrates in the cell including ribosomal RNA and those that directly or indirectly affect the mRNA levels of dozens of genes, and thus presumably affect their cognate protein expression levels (Sim, et al. 2010; Stead, et al. 2011; Gordon, et al. 2017). Mutations in RNase III thus have the potential to affect fitness through a variety of mechanisms. We developed our fluorescence-based, high-throughput functional score to interrogate the relationship between RNase III function and fitness. However, our functional score reports on the cleavage of just one RNase III substrate, the 5’ UTR of its own mRNA. Despite using a single substrate, our functional score and fluorescence assay can explain most of the variation in fitness among our selected mutants (**fig. 4D,E**), and a positive correlation between functional score and fitness among all mutations is apparent (**fig. 3E**). This suggests that the 5’UTR of *rnc* is a good, representative RNase III substrate. Mutational changes in the rate of cleavage of the 5’UTR of *rnc* reflect changes in the rate of cleavage of the RNase III substrate(s) that impact fitness. Such an outcome with our chosen substrate was not guaranteed, as one can imagine specificity-altering mutations that differentially affect the cleavage of the 5’UTR of *rnc* and RNase III substrates important for fitness.

Visual comparison of the fitness, transformation, and functional landscapes (**fig. 2**) does not suggest that specificity-altering mutations are common, though limitations in the resolution of our measurements make it difficult to discover single substitutions that alter specificity or positions with small effects on RNase III specificity. However, our data suggests F188 is important for RNase III specificity. The fitness and transformation score effects of F188D were much more deleterious than expected based on all three functional measures (**fig. 4D-F, H**). F188 is highly conserved and lies in a *β*-strand right after RBM2 (residues 181-186) only ∼3.4 Å from the dsRNA in our structural model.

Interestingly, no amino acid substitution at F188 had a significantly deleterious functional score, yet several substitutions at F188 are deleterious for both fitness and the transformation score, most notably mutations to Asp, Glu, Pro, and Gly (**fig. 4G,H**), which are four of the five least common amino acids in *β*-strands (Fujiwara, et al. 2012). This difference suggests that these F188 mutations, while not being very deleterious for cleavage of the 5’UTR of *rnc*, are more deleterious for cleavage of at least one other RNase III substrate, specifically one that is consequential for fitness.

### RNase III’s tolerance to mutation

A common way to quantify the sequence conservation of a protein family is through calculation of the Shannon entropy at each residue using a multiple sequence alignment of the family members (Shenkin, et al. 1991). The calculation makes use of the probability of finding each amino acid at that position, values which vary from 0 to 1. From this entropy each position’s *k\** value can be calculated, which is considered the average number of amino acids that are tolerated at a position. A *k\** of 20 means that all 20 amino acids are equally probable, while a *k\** of 1 means that only one amino acid is found at that position. These *k\** values are a prediction of the mutational tolerance by position for the protein family based on what is observed naturally.

We have proposed an analogous calculation for position-dependent mutational tolerance of a single protein using the set of fitness values from DMS experiments, provided the fitness values range from 0 to 1 (Firnberg, et al. 2014). These *k\** values are a measure of the actual mutational tolerance at each position for that protein. For RNase III, fitness values do not range from 0 to 1, but the relative enrichment values used to calculate the transformation and functional scores nearly do. Thus, using the relative enrichment values we calculated *k\** values for transformation and functional scores. We compared these values to each other and to *k\** values we determined from a multiple sequence alignment of 355 unique UniProt-curated RNase III family sequences (**fig. 5**). The distribution of *k\** values indicate that many positions in *E. coli* RNase III have a high degree of mutational tolerance, with more than 50% tolerating 15 or more different amino acids (**fig. 5A,B**). We find that if a position is relatively variable in the RNase III family, the position is almost always highly tolerant to mutation in the *E. coli* protein (**fig. 5C**). However, the inverse is not necessarily true. While key active site residues have low *k*\* values in all calculations, several positions that are largely invariant in nature can tolerate many mutations in the *E. coli* protein.

**Figure 5.**
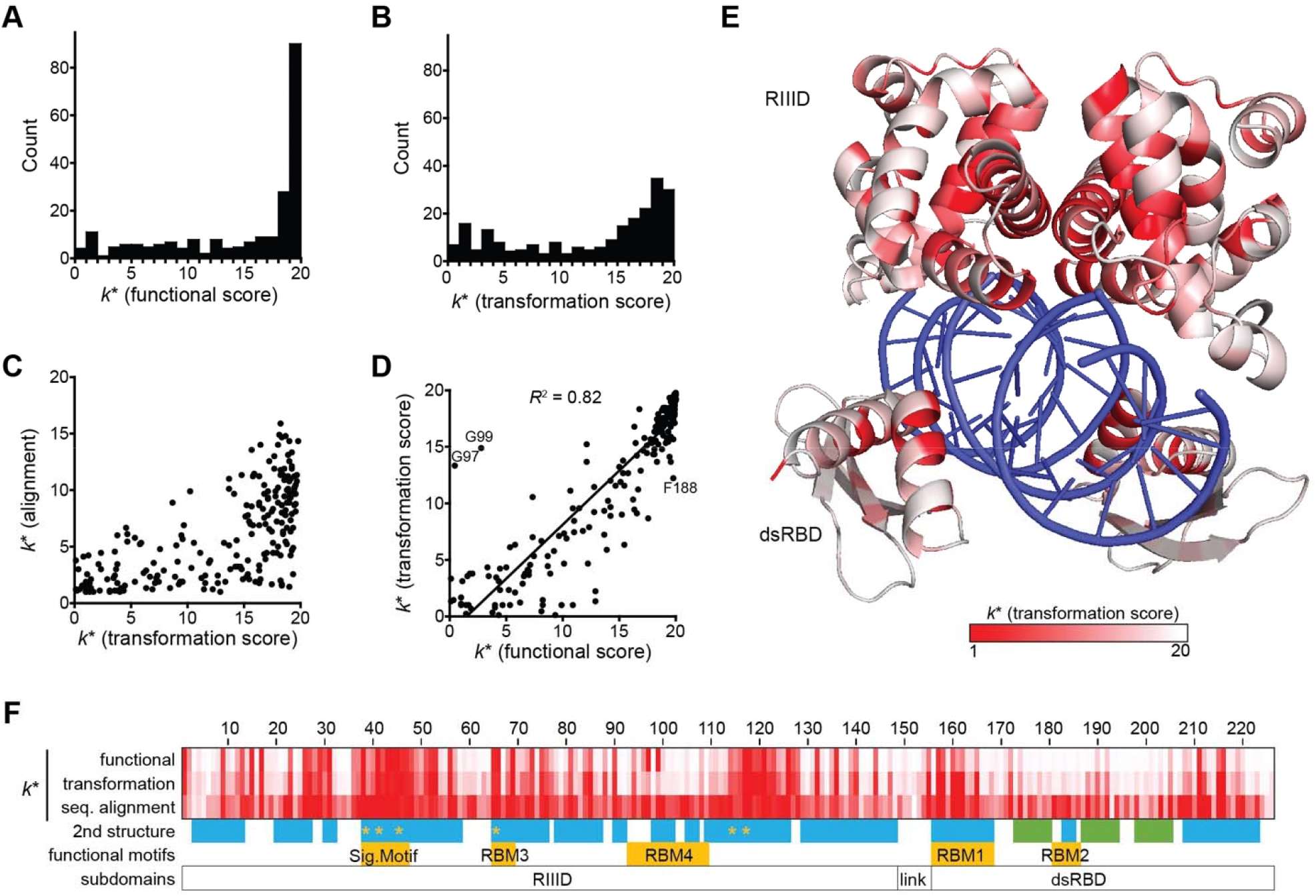
RNase III mutational tolerance. Mutational tolerance as show by the distribution of *k*\* values for (**A**) functional score and (**B**) transformation score values. (**C**) Correlation of transformation score *k*\* values with those from a sequence alignment of the RNase III family. (**D**) The *k*\* values for functional score and transformation score correlate linearly. (**E**) Transformation score *k*\* values mapped onto a structural model of *E. coli* RNase III bound to an dsRNA substrate. RNase III model was generated from homology modeling to the *T. maritima* (1O0W) and *A. aeolicus* (2NUG) RNase III structures using Swiss Prot. The linker between the RIIID and the dsRBD is not included as it was poorly resolved in the 1O0W and 2NUG structures. (**F**) A comparison of *k*\* values mapped on the primary and secondary structure of *E. coli* RNase III. The range of *k*\* is 1 (white) to (20) red for functional and transformation scores and 1 (white) to 15 (red) for the sequence alignment. All *k*\* values are tabulated as **supplementary data S4**, Supplementary Material online.

We find that *k*\* values calculated from transformation and functional scores correlate linearly (*R*^2^ = 0.82), further illustrating how catalytic activity is the major determinant of fitness (**fig. 5D**). Interestingly, positions G97 and G99 in RBM4 were strong outliers of this trend. Except for mutations to small residues, which are near neutral for function and fitness at this position, most mutations at G97 and G99 cause a much smaller reduction in transformation scores than would be predicted from their large deleterious effect on functional scores (**fig. 5D**). This result indicates that although cleavage of the 5’UTR of *rnc* strongly requires small amino acids at G97 and G99, cleavage of RNase III substrates that are important for fitness are not nearly so negatively impacted by mutations to large residues at these positions despite both G97 and G99 being highly conserved in the RNase III family (*k*\* values 1.9 and 2.1, respectively). This result predicts that mutations to non-small amino acids at G97 and G99 substantially impact the sequence specificity of RNase III.

### RNase III fitness and functional landscapes in the context RNase III’s known sequence-structure-function relationship

The RNase III monomer can be divided into three regions: the RIIID, dsRBD, and the flexible linker that connects these two regions. RNase III’s positional tolerance to mutation, as measured by *k*\* values for transformation score, are broadly consistent with the structural model of RNase III (**fig. 5E**). Positions less tolerant to mutation tend to fall at the protein-RNA and dimer interfaces or in the interior of the protein, with surfaces residues tending to be more tolerant. Within the RIIID are all the active site residues responsible for cleavage of dsRNA as well as all the points of contact between the two monomers. Each RNase III active site is composed of six negatively charged residues that contribute to positioning and cleavage of the dsRNA: E38, E41, D45, E65, D114, and E117 (Blaszczyk, et al. 2001; Gan, et al. 2006; Gan, et al. 2008; Court, et al. 2013). We find very low tolerance for mutations at these positions except for D114 (**fig. 5F** and **supplementary fig. S9**, Supplementary Material online). At positions E41, D45, and E117, no mutation is tolerated confirming previous reports that these residues are essential for RNase III activity (Inada and Nakamura 1995; Dasgupta, et al. 1998; Blaszczyk, et al. 2001; Sun and Nicholson 2001; Sun, et al. 2004; Zhang, et al. 2004). Three of these residues are in RNase III’s signature motif of ten residues (residues 38-47). The motif is highly conserved across the RNase III family of proteins (Mian 1997; MacRae and Doudna 2007; Court, et al. 2013; Nicholson 2014) and generally intolerant to mutations in this study (**fig. 5F** and **supplementary fig. S9**, Supplementary Material online).

Assisting the six negatively charged active site residues are two RNA binding motifs (RBM) in the RIIID that help position the scissile bond at the correct positions for cleavage (Court, et al. 2013). RBM3 shows little tolerance to mutations except to like residues and those found in RNase III family enzymes in other organisms. Binding occurs between RBM3 and dsRNA when S69 forms a hydrogen bond with the RNA at the cleavage site and the side chain of E65 forms a hydrogen bond within the dsRNA major groove (Gan, et al. 2006). Most mutations at E65 are very deleterious, but quite surprisingly E65P is neutral for fitness and among the least deleterious for the functional score. RBM4 was much more tolerant to mutations than RBM3 (**fig. 5F**; median values 0.99±0.01 versus 0.90±0.11, *p*< 0.0001, Wilcoxon rank-sum test). RMB3’s greater sensitivity to mutation is consistent with its active site proximity and greater number of contacts with the dsRNA. The most severe mutations in RBM4 were those to G97 and G99, but only for the functional score as described previously.

The dsRBD was more robust to mutations than the RIIID in each of the landscapes. The dsRBD is composed of two α-helices and a β-sheet made up of three antiparallel strands. Most of the deleterious mutations in the dsRBD were within the two α-helices, whereas the β-sheets were less affected by mutation. The two RNA binding motifs within the dsRBD are found in the first α-helix (RBM1) of the domain and the loop connecting the β1 and β2 strands (RBM2). RBM1 was much less tolerant to mutation than RBM2 (**fig. 5F**), as might be expected (Gan, et al. 2006) due to its cleavage site proximity and larger number of dsRNA interactions. Residues in RBM2 interact with two bases within the minor groove of the RNA and the side chain of H184 makes one hydrogen bond with an O2’ hydroxyl (Gan, et al. 2006) but these interactions must be of minor importance for function and fitness based on RBM2’s high mutational tolerance.

The dsRBD is essential for activity in vivo (Nicholson 1999; Sun, et al. 2001), although RNase III lacking the dsRBD is capable of dsRNA-specific cleavage in vitro in non-physiological salt concentrations (Sun, et al. 2001). Our DMS data indicated that the dsRBD is essential as the region was intolerant to nonsense mutations, except for the final three residues. To confirm this result, we created a ΔdsRBD mutant through triple mutation of Q153*/K154*/D155* to test the effects of removing the dsRBD (**Table 1**). We found that deleting the dsRBD reduced fitness to 0.75 ± 0.01, equivalent to cells lacking RNase III. The fluorescent assay and cleavage assay indicated Q153*/K154*/D155* was severely impaired in cleaving the *rnc* 5’UTR, thus confirming that the dsRBD is required for in vivo cleavage activity.

The flexible linker region connecting the RIIID and dsRBD (residues 149 – 155) was fairly tolerant to mutation except for D155, the most highly conserved linker positions (**fig. 5F**). Within the linker, mutations to proline, especially at G150, were the most deleterious. Based on structural models of *T. maritima* RNase III with and without dsRNA, free rotation is observed around the flexible linker until both the RIIID and dsRBD have complexed with the dsRNA (Gan, et al. 2005, 2006). Introduction of a rigid proline with the linker, particularly at G150, likely interferes with the linker adopting the necessary conformation to orient the two domains correctly.

## Conclusions

Our study represents the first comprehensive report on both the fitness and in vivo functional effects of missense mutants to the same gene. We developed three landscapes each reporting on different aspects of the effect of RNase III mutations. Our fitness landscape shows how a mutation affects the growth rate of cells during exponential growth in liquid media, whereas the transformation score fitness landscape shows how each mutation affects the fitness of cells during plasmid transformation and subsequent growth on solid media. The functional landscape shows how each mutation affects the cellular level of RNase III catalytic activity on its own 5’UTR. We saw that both fitness and transformation scores strongly correlate with functional scores, indicating that the ability of RNase III to properly cleave dsRNA is the major fitness determinant for this gene.

The distribution of fitness and functional effects were both bimodal, with the largest fraction of mutations near neutral, but a sizable fraction of mutations causing a fitness effect approximately equivalent to that of loss of RNase III function. We observed that key residues including the signature motif, active sites residues, RBM1, and RBM3 were largely intolerant to mutation in each of our landscapes. A comparison of the RNase III family’s sequence diversity and *E. coli* RNase III’s mutational tolerance (**fig. 5**) showed that positions in *E. coli* RNase III that were intolerant to mutation lacked sequence diversity in the RNase III family. Conversely, however, some regions that lacked diversity in nature were highly tolerant to mutation in *E. coli* RNase III, most notably RBM4 and RBM2 and the *β*-strands in the dsRBD. We identified three residues (G97, G99, and F188) at which mutations have significantly different effects on functional score and fitness. These differences are likely due to the mutations having different effects on the cleavage of the 5’UTR of *rnc* and the cleavage of RNase III substrate(s) important for fitness (i.e. the mutations impact RNase III substrate specificity).

## Materials and Methods

### *E. coli* strains and growth conditions

Strain NEB5α (*fhu2 (argF-lacZ)U169 phoA glnV44 80 (lacZ)M15 gyrA96 recA1 relA1 endA1 thi-1 hsdR17)*(C2987) was purchased from New England Biolabs. Strain MG1693 (*E. coli* MG1655: K12 F^−^ lambda^−^ *ilvG*^−^ *rfb*-50 *rph*-1 *thyA715)* and SK7622 (MG1693 Δ*rnc-38* Km^R^) were a gift from Sidney Kushner. The PH0919 strain was a gift from Pricila Hauk.

Except where noted, all LB media was supplemented with 50 µg/mL thymine, 100 µg/mL ampicillin to maintain pRnc2-gRNA, and 50 µg/mL chloramphenicol to maintain pdCas9. Overnight cultures were grown in test tubes containing 10 mL LB-media supplemented with appropriate antibiotics shaking at 250 rpm at 37°C for 16 h. Overnight cultures of *rnc* ^*–*^ strains were grown overnight supplemented with 25 µg/mL kanamycin and transferred to media without kanamycin for experimentation. Growth experiments were performed in 500 mL baffled shake flasks containing 100 mL LB-media supplemented as above and with 0.2% w/v glucose shaking at 250 rpm at 37°C.

### Vectors

pRnc2-gRNA (#Addgene #189563) was made by inserting the Phusion High Fidelity polymerase (New England Biolabs) amplified *rnc* 5’-UTR (nucleotides 1-160) in place of the pBAD 5’-UTR on pRnc-gGFP129 (Hauk, et al. 2022) by blunt ligation. pdCas9 (#44249) was purchased from Addgene.

### Monoculture growth assay for *rnc* ^−^ growth rate

MG1693 and SK7622 (*rnc* ^*-*^) cells harboring pRnc2-gGFP, pRnc2-GFP with mutant *rnc* alleles, or the empty vector (EV) were grown overnight and diluted to an (optical density) OD_600_ of 0.05 in unbaffled shaker flasks. OD_600_ was monitored until cells reached the exponential growth phase (∼OD_600_ of 0.5) and then diluted to an OD_600_ of 0.02 in baffled shaker flasks containing pre-warmed to 37°C LB-media. Cells were grown for approximately 10 generations; OD_600_ was measured for every 30 min for approximately 6 h with a 3.1 ml/96.9 mL dilution in new pre-warmed to 37°C LB-media at about 3 h.

Fitness was determined as the ratio of the mean growth rate of cells expressing the *rnc* mutant to the mean growth rate of cells expressing the wildtype *rnc*. The mean growth rate was calculated from the OD_600_ data as a function of time between the first time point (30 min) and final timepoint (after ∼ 10 generations of growth). The growth rate was the slope of the linear fit of ln(OD_600_) vs. time.

### Library creation

Libraries of all possible single-codon substitutions in *rnc* were constructed essentially as described (Mehlhoff, et al. 2020); the protocol is described in detail here. All single-codon substitutions were generated by inverse PCR with Phusion High Fidelity polymerase in HF Buffer (New England Biolabs) using degenerative codon NNN targeted to each codon of the *rnc* gene. Amplification at each codon was verified by gel electrophoresis on a 1% TAE-agarose gel. Inverse PCR and gel verification was repeated for any codons lacking prominent bands corresponding to the size of pRNc2-gRNA (4.8 kb). Successfully amplified vectors were pooled together based on the relative intensity of bands. Plasmids were pooled in three sublibraries – one for each third of the gene (region 1, residues 1-75; region 2, residues 76-150,; region 3, residues 151-226). Sublibraries were used due to read constraints of Illumina MiSeq. Pooled DNA was separated on a 1% TAE-agarose gel and desired band extracted using PureLink Quick Gel Extraction Kit (Invitrogen). The extracted DNA was cleaned and concentrated using a DNA Clean & Concentrator Kit (Zymo) and phosphorylated using T4 polynucleotide Kinase (New England Biolabs). The phosphorylated DNA was cleaned and concentrated again as above and then ligated using T4 DNA Ligase (New England Biolabs). The ligation products were transformed into NEB5α electrocompetent cells and plated on LB-agar bioassay dishes supplemented with 50 µg/mL ampicillin and 2% w/v glucose. We estimated that a target of 22,000 transformants was sufficient to constitute a library that was about 99% complete (Bosley and Ostermeier 2005) and repeated the transformation step until the number of transformants exceeded 22,000 in total. Successful library creation was verified by Sanger sequencing of random transformants (Genewiz). All colonies were collected from bioassay dishe in 15 mL of LB-media containing 15% glycerol, centrifuged at 3000 xg, and the pellet was resuspended in a small volume of supernatant then stored at -70°C.

An aliquot of each region of the NEB5α *rnc* library was thawed on ice and DNA purified using a QIAprep Spin Miniprep Kit (Qiagen). DNA concentration was estimated by NanoDrop spectrophotometer and verified by gel electrophoresis on a 1% TAE-agarose gel. To avoid double mutant transformants (Goldsmith, et al. 2007), 8.3 ng of each region was transformed into electrocompetent SK7622 cells for the fitness landscape experiment and electrocompetent PH0919 cells harboring the pdCas9 plasmid for the functional landscape experiment. Colonies were collected from the transformation plates as before, centrifuged, resuspended in a small volume of supernatant, and stored at -70°C.

### Growth competition experiment of the SK7622 library

Growth competition was performed essentially as described (Mehlhoff, et al. 2020). SK7622 frozen library stocks for each of the three regions were diluted to OD_600_ of 0.01 in 100 mL of LB-media in unbaffled shaker flasks and incubated until the OD_600_ reached approximately 0.5. Ten mL of this culture was placed on ice and centrifuged at 3000 xg for 10 min at 4°C. The cultured cells at OD_600_ of 0.5 were diluted to OD_600_ of 0.02 in 100 mL of prewarmed to 37°C LB-media in baffled shake flasks. Flasks were incubated at 37°C with shaking until OD_600_ was approximately 0.640 (5 generations of growth). The culture was diluted at a ratio of 3.1 mL culture to 96.9 mL of new pre-warmed to 37°C LB-media in a separate baffled shaker flask and incubated at 37°C until OD_600_ reached approximately 0.64 (5 additional generations of growth). Ten mL of this culture was placed on ice and centrifuged at 3000 xg for 10 min at 4°C. Plasmid from the initial and final time point was extracted and purified from pelleted cells using a QIAprep Spin Miniprep Kit (Qiagen).

### Transformation score experiment

Frozen NEB5α and SK7622 library stocks were thawed on ice then centrifuged at 3000 xg for 10 min at 4°C. Plasmid DNA was extracted from pelleted cells. Extracted DNA was stored then prepared for Illumina MiSeq as described below.

### Functional score experiment

Electrocompetent PH0919 cells containing pdCas9 were transformed with pRnc2-gGFP with wildtype RNase III or the E117K mutant, incubated overnight on bioassay dishes, and colonies collected then frozen as described above. Aliquots of the frozen PH0919 library, wildtype, and E117K mutant were thawed on ice. Twenty µL of frozen cells were diluted into 1980 µL of cold phosphate-buffered saline (PBS) and the OD_600_ was measured. Cells were then diluted to an OD_600_ of 0.1 in a final volume of 2 mL; region 2 samples were spiked with 1% E117K sample based upon OD_600_. Fluorescence-activated cell sorting (FACS) of each library region was performed based on GFP-fluorescence and forward scatter gating placed around wildtype RNase III and E117K samples using a Becton Coulter MoFlo Legacy cell sorter. Cells were sorted until greater than 1 × 10^6^ high fluorescent events were collected, during which time approximately 1.5-2.0 × 10^7^ low fluorescent events were collected. Sorted cells were centrifuged at 3000xg and DNA extracted, stored, and then prepared for Illumina MiSeq as described below.

### Illumina MiSeq sample preparation

Plasmids collected during the growth competition experiment, the transformation score experiment and the functional score experiment were digested with XhoI (New England Biolabs) and purified using a DNA Clean & Concentrator Kit (Zymo). Custom Illumina adapter sequences (IDT) containing unique barcode to identify which region and time point DNA corresponded to were added to the linearized DNA by PCR amplification with Phusion High Fidelity polymerase (New England Biolabs) and HF buffer (New England Biolabs). For replica 1 of the growth competition experiment, PCR amplification was performed with ∼50 ng of DNA template and following a protocol of 98°C for 30 sec, 25 cycles of 98°C for 15 sec, 54°C for 15 sec, and 72°C for 1 min, followed by 72°C for 5 min. For replica 2 of the growth competition experiment and the transformation experiment, PCR amplification was performed as replica 1 but with ∼25 ng of DNA template and following a protocol with 15 cycles instead of 25. For replica 1 and 2 of the functional experiment, PCR amplification was performed as above with ∼25 ng of DNA and following a protocol with 20 cycles instead of 25. Proper DNA size was verified by gel electrophoresis on a 2% TAE-agarose gel. Samples were pooled and submitted for Illumina MiSeq (2 × 300 bp reads) at the Transcriptomics and Deep Sequencing Core facility at Johns Hopkins University.

### Deep sequencing analysis

Illumina MiSeq reads were analyzed essentially as described (Mehlhoff, et al. 2020). Paired-end reads were merged using PEAR (Zhang, et al. 2013) set to a minimum assembly length of 200 base pairs. Illumina adapters and base pairs outside the desired region were cropped from the merged reads using Trimmomatic (Bolger, et al. 2014) (Region 1-Headcrop: 39, crop 225; Region 2-Headcrop: 33, crop: 225; Region 3 – Headcrop: 35, crop: 228). The resulting trimmed read were input to Enrich2 (Rubin, et al. 2017), which counted the variants for use in calculating enrichment to determine fitness, transformation score, functional score, and corresponding variances. Reads containing bases with a quality score below 20, bases marked as N, or mutations at more than one codon were filtered out.

### Fitness, transformation score, and functional score calculation

Fitness values (*w*) were determined as described (Mehlhoff, et al. 2020) using Equation 1, except the rate of wildtype synonyms was used in place of the growth rate for wildtype

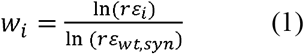

where *r* is the fold increase in total number of cells and ε_*i*_ is the enrichment of non-synonymous mutation *i*. Fitness was calculated on the amino acid level by pooling the sequencing counts for all synonymous codons.

Enrichment of non-synonymous mutations was calculated as described (Mehlhoff, et al. 2020) using Equation 2 where c_*i*_ is the count of non-synonymous mutation *i* in the designated population

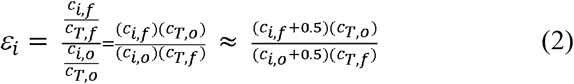

and C_*T*_ is the total count of the designated population. C_i_ was calculated by summing the counts of all synonymous mutations for each amino acid substitution. We added the 0.5 to all counts of mutants to assist with mutations that have counts that are zero (i.e. so that a fitness value can still be calculated).

Since there was not cell growth in the transformation and functional experiments, we defined the transformation and functional scores as:

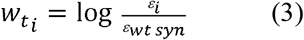

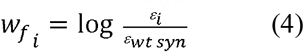

The enrichments were determined using Equation 2. For the transformation score, the NEB5α library was population *o*, while the SK7622 library was population *f*. For the functional score, the highly fluorescent (i.e. most similar to E117K) population was population *o* while the dim GFP population (most similar to wildtype) was population *f*.

### Statistical treatment of DMS measurements

The statistical analysis for fitness values was performed as described (Mehlhoff, et al. 2020). Statistical calculations for transformation and functional scores were calculated as below. For transformation score, the NEB5α library was population *o*, while the SK7622 library was population *f*. For the functional score, the highly fluorescent (i.e. most like E117K) population was population *o* while the dim GFP population (most like wildtype) was population *f*. The pre-log variance of transformation score and functional score was defined as the variance *σ*^*2*^, which depends on the proportion *p* and the number of samples from the population:

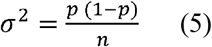

For our calculations 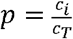 and n = c_*T*_.

In propagating uncertainty, the variance of a proportion of proportions was defined as:

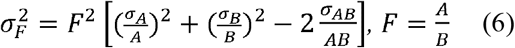

Here the 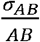 term was equal to 0 since there is no covariance between proportions A and B. To calculate the variance for transformation score and functional score, there were four proportions to consider: the frequencies of the mutant in the *o* and *f* populations, and the frequency of WT synonyms in *o* and *f* populations. Thus, the variance of the functional fitness was defined as :

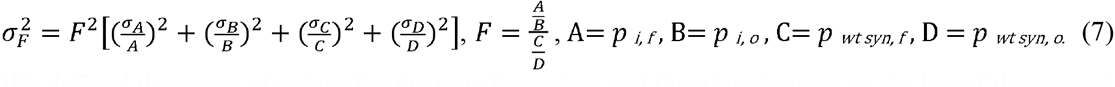

We defined the range of values for the transformation and functional scores as the log of the score ± variance from equation 7.

Confidence intervals and *p*-values for transformation and functional scores were calculated as described (Mehlhoff, et al. 2020) using the variance.

### RNase III fluorescence activity assay

Overnight inoculums were diluted to a final OD_600_ of 0.00625 in 100 mL of LB-media in baffled shake flasks. Cultures were incubated at 37°C with shaking at 250 rpm for approximately 4 hours. Triplicates of 100 µL from each culture were added to a clear/black bottom 96-well microplate. OD_600_ and GFP fluorescence at 485 nm excitation/525 nm emission were measured using a SpectraMax M3 microplate reader.

### In vitro primer extension

The primer extension assay was performed essentially as described (Sharma and Woodson 2020); the protocol is described in detail here (Hauk, et al. 2022). After 6 hr of growth, 1 mL of 1x 10^8^ cells/mL were harvested and pelleted at 3000x g. The cell pellet was incubated in 200 µL of TE buffer (Gentrox) containing 1 mg/mL lysozyme at room temperature for 5 min, vortexing for 10 s every 90 s. A total of 680 µL of RLT (Qiagen) buffer with 1% 2-mercaptoethanol was added to the resuspended pellet followed by addition of 490 µL of 100% ethanol. This lysate was used for RNA extraction with an RNeasy Mini Kit (Qiagen) according to the manufacturer’s standard protocol for bacterial RNA purification.

The purified RNA was used to amplify cDNA from the gUTRGFP template with the 5’ Cy-5 labelled pWP252fluor primer (5’ ATAACGGACTAGCCTTA 3’) using the standard Superscript III First-Strand Synthesis System (Thermo Fisher) with 3 µg of total RNA and elongating at 52.5°C for 60 min. Next, undyed urea loading buffer (8 M urea, TE buffer) was added to cDNA and the DNA was denatured at 95°C for 3 min prior to loading in Novex 6% TBE-urea gel (Thermo Fisher). Samples were separated for 45 min at 180 V, 19 mA in TBE buffer (Quality Biological). The gel was visualized for Cy-5 fluorescence using a Typhoon 9410. DNA standards (ssDNA standards) were amplified from pRnc-gRNA DNA using Phusion High Fidelity DNA Polymerase (New England BioLabs) with the pWP252fluor reverse primer and forward primers annealing to the expected 5’ termini of the cleaved and uncleaved 5’UTR that would yield the desired size products. PCR products were separated on a 4% agarose gel at 110V for 40 min and proper-sized products extracted using QIAquick gel extraction kit (Qiagen). Following denaturation in undyed loading buffer at 95°C for 3 min, DNA standards and cDNA both migrate as ssDNA in denaturing gels.

Cleavage scores were quantified using ImageJ software (Schneider, et al. 2012). Cleavage percentages are determined as the ratio of the intensity of the sum of upstream and downstream cleavage bands to the sum of the uncleaved, upstream and downstream cleaved bands. Cleavage scores represent the mean of three biological replicates each with two technical duplicates.

## Acknowledgments

We thank Hao Zhang for his support with FACS and Jacob Mehlhoff for his assistance with deep sequencing analysis. This research was supported by National Science Foundation grants MCB-1817646 and MCB-2113019 to M.O.

## Data availability

DMS sequencing data can be found at BioProject (https://www.ncbi.nlm.nih.gov/bioproject) using accession no. PRJNA887445. Sequencing counts for the DMS studies can be found in **supplementary data S1-3**, Supplementary Material online.

## Supplementary Materials for

### This PDF file includes

Figs. S1 to S9

Captions for Data S1 to S4

### Other Supplementary Materials for this manuscript include the following

Data S1 to S4

**Supplementary Figure S1:**
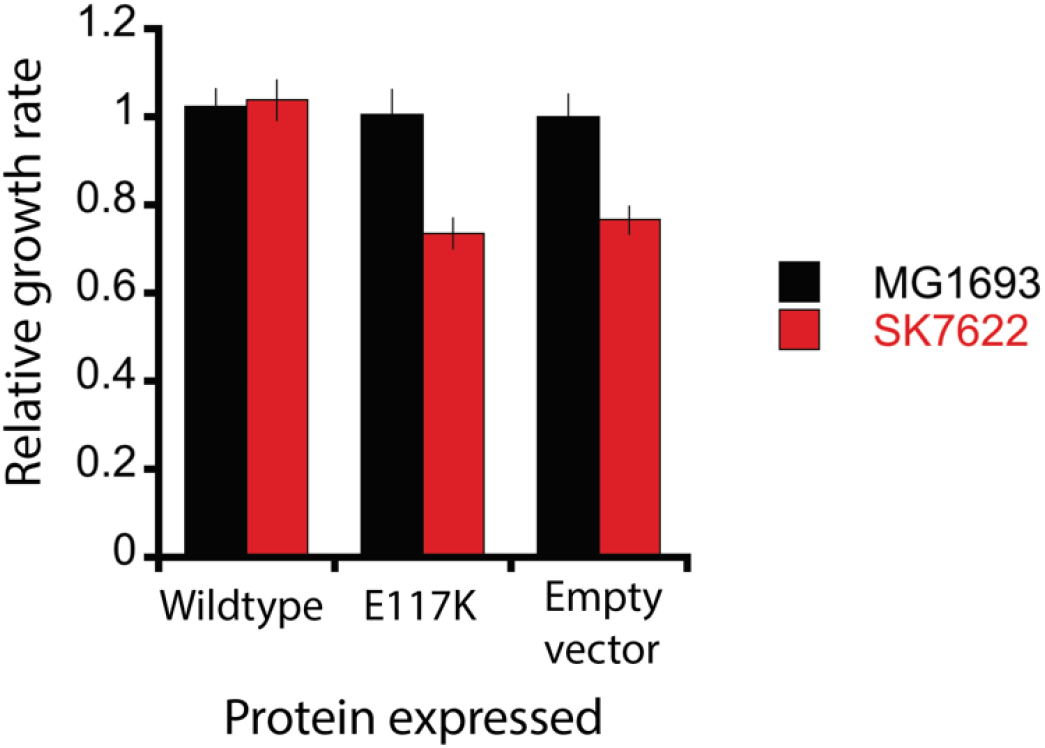
E. coli cells lacking a functional RNase III grow 25% slower. MG1693 and SK7622 (MG1693 *rnc*^*–*^) cells harboring pRnc2-gGFP plasmids expressing the wildtype RNase III, nuclease-null RNase III E117K variant, and an empty vector were grown in LB with 0.2% glucose for 10 generations. Values are the mean relative growth rate normalized to that of MG1693 harboring the empty vector. Error bars represent the standard deviation (*n* = 4).

**Supplementary Figure S2:**
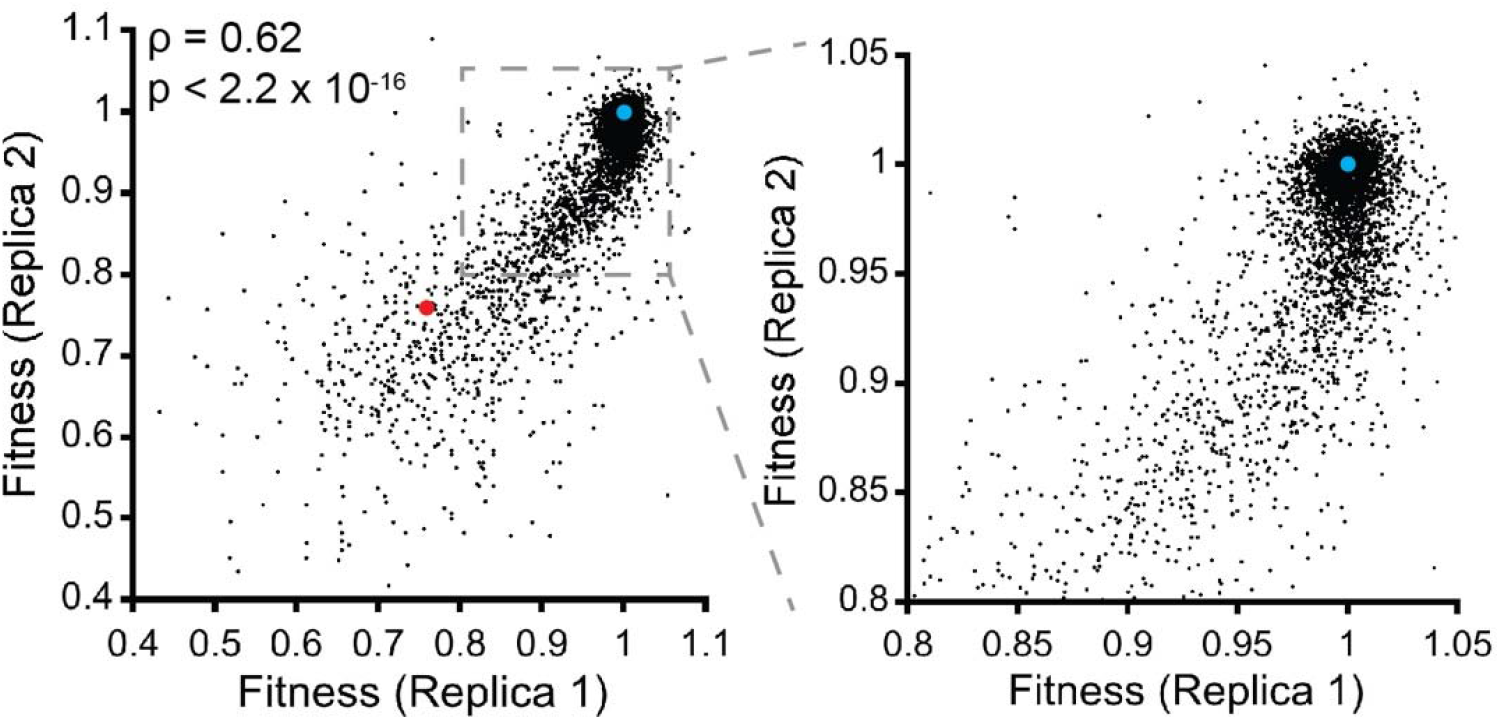
Correlation between fitness effects observed in replica 1 and replica 2. The blue point represents wildtype, and the red point represents nonfunctional RNase III as measured by the growth rate of cells with the M1S mutation (i.e. start codon replaced with TCT) relative to that of wildtype.

**Supplementary Figure S3:**
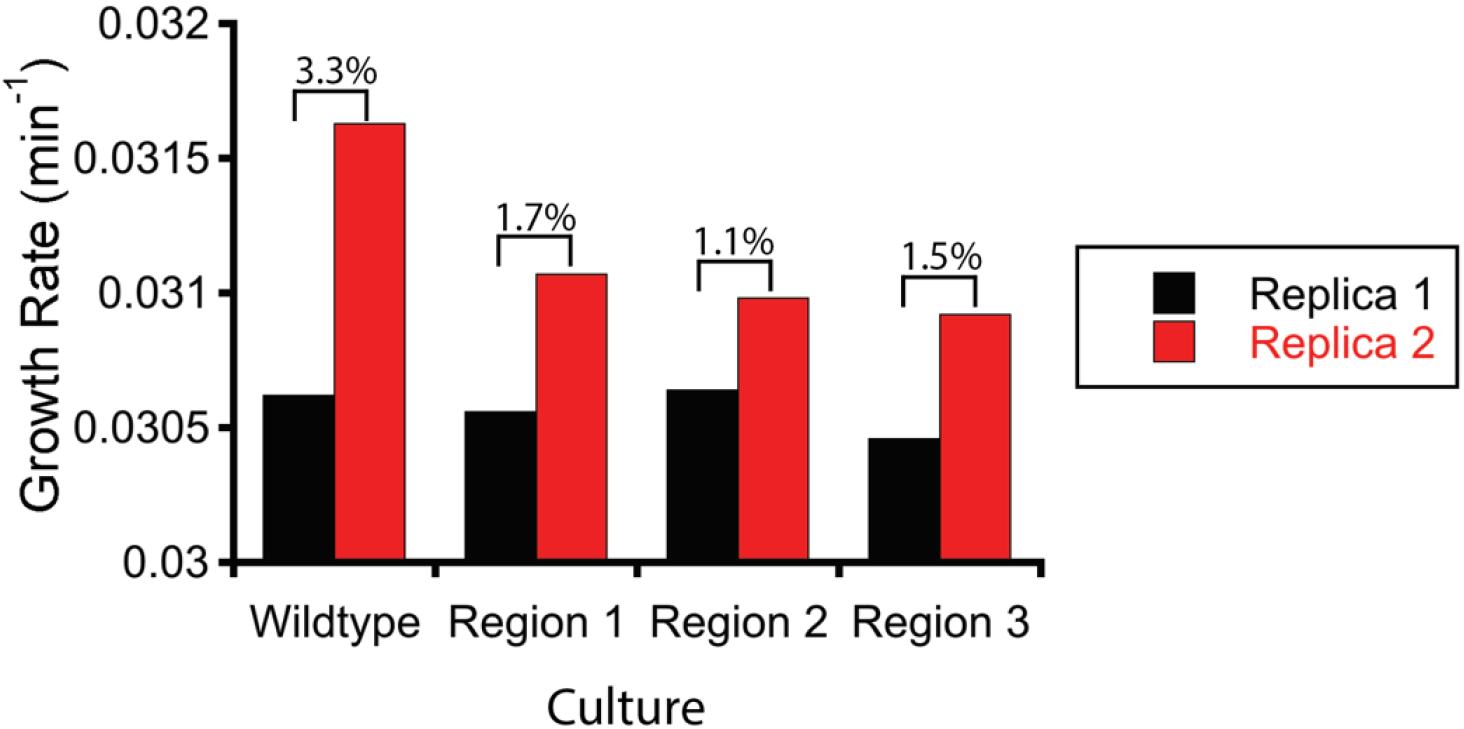
Absolute growth rates during growth competition experiments to determine fitness values. The percentages indicate how much faster replica 2 was over replica 1.

**Supplementary Figure S4:**
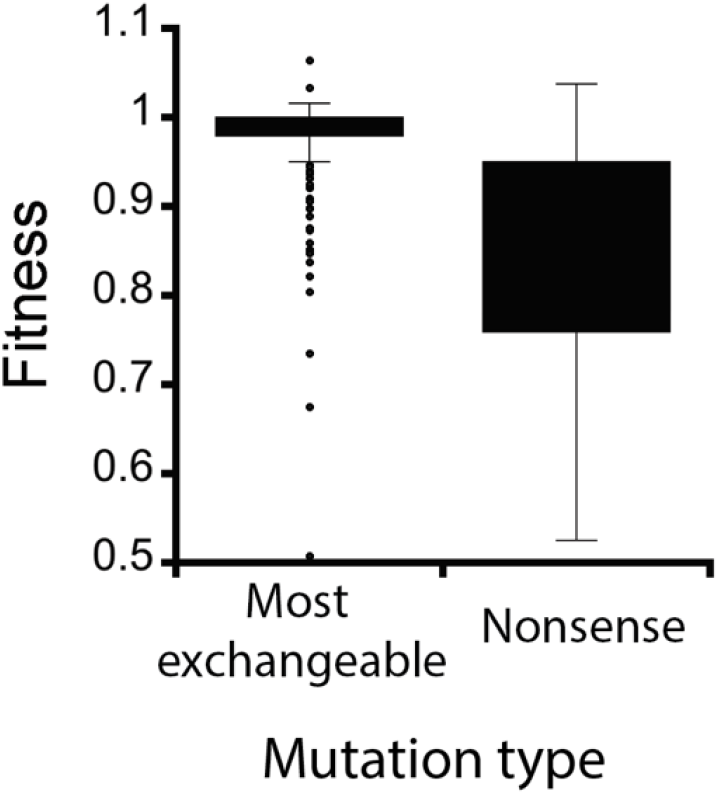
Nonsense mutations and conservative mutations have very different fitness value distributions. Conservative mutations, defined as mutations to the most exchangeable amino acid (Yampolsky and Stoltzfus 2005), show a much narrower range of fitness values than nonsense mutations. Nonsense mutations were more deleterious than conservative mutations (*p*-value <0.0001 by Welch’s T test).

**Supplementary Figure S5:**
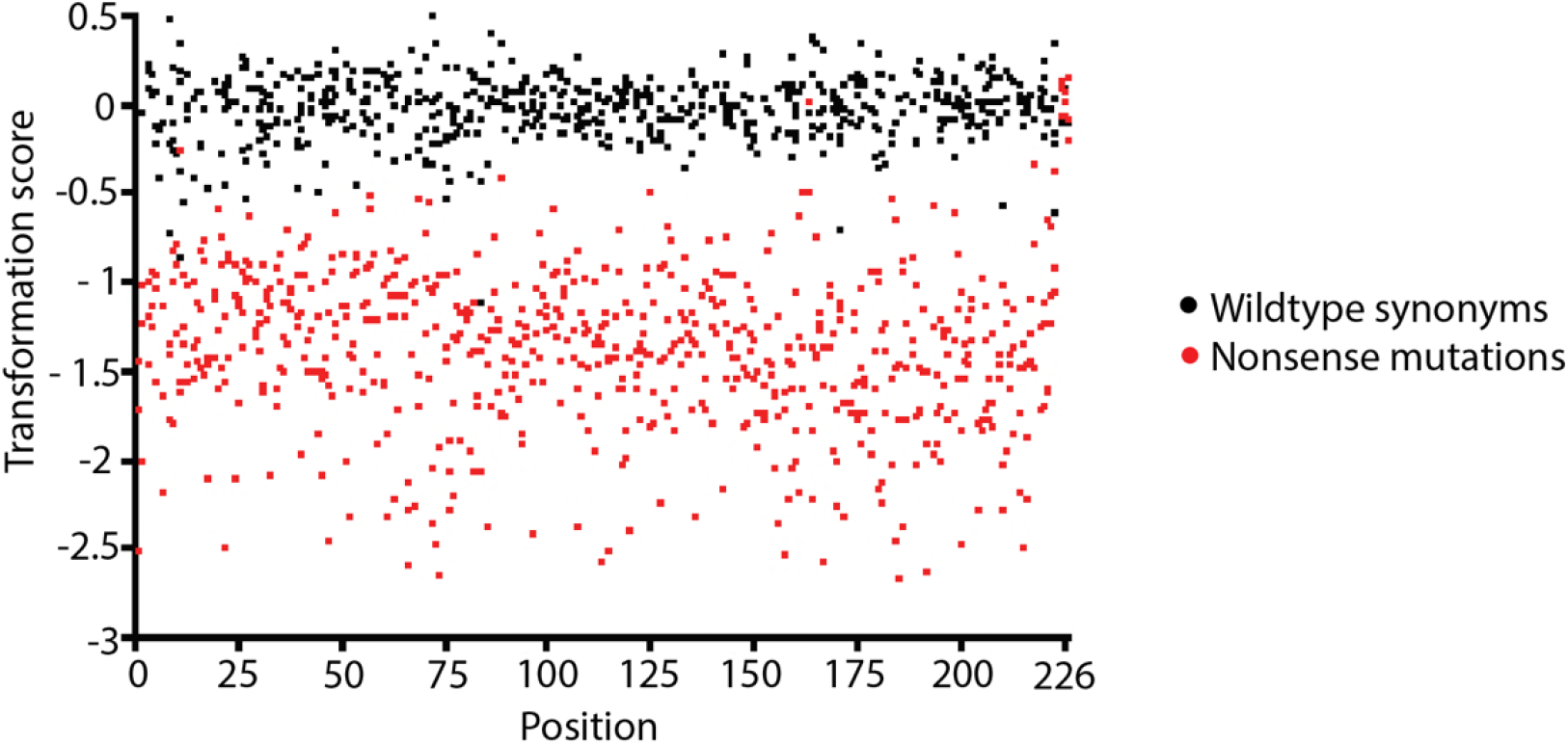
Transformation into a *rnc*^−^ strain depletes nonsense mutations but not mutations synonymous to wildtype. Transformation scores for wildtype synonyms and nonsense mutations at each position within RNase III. The weighted mean transformation scores for wildtype synonyms and nonsense mutations (residues 1-223) are 0.00 and -1.36, respectfully.

**Supplementary Figure S6:**
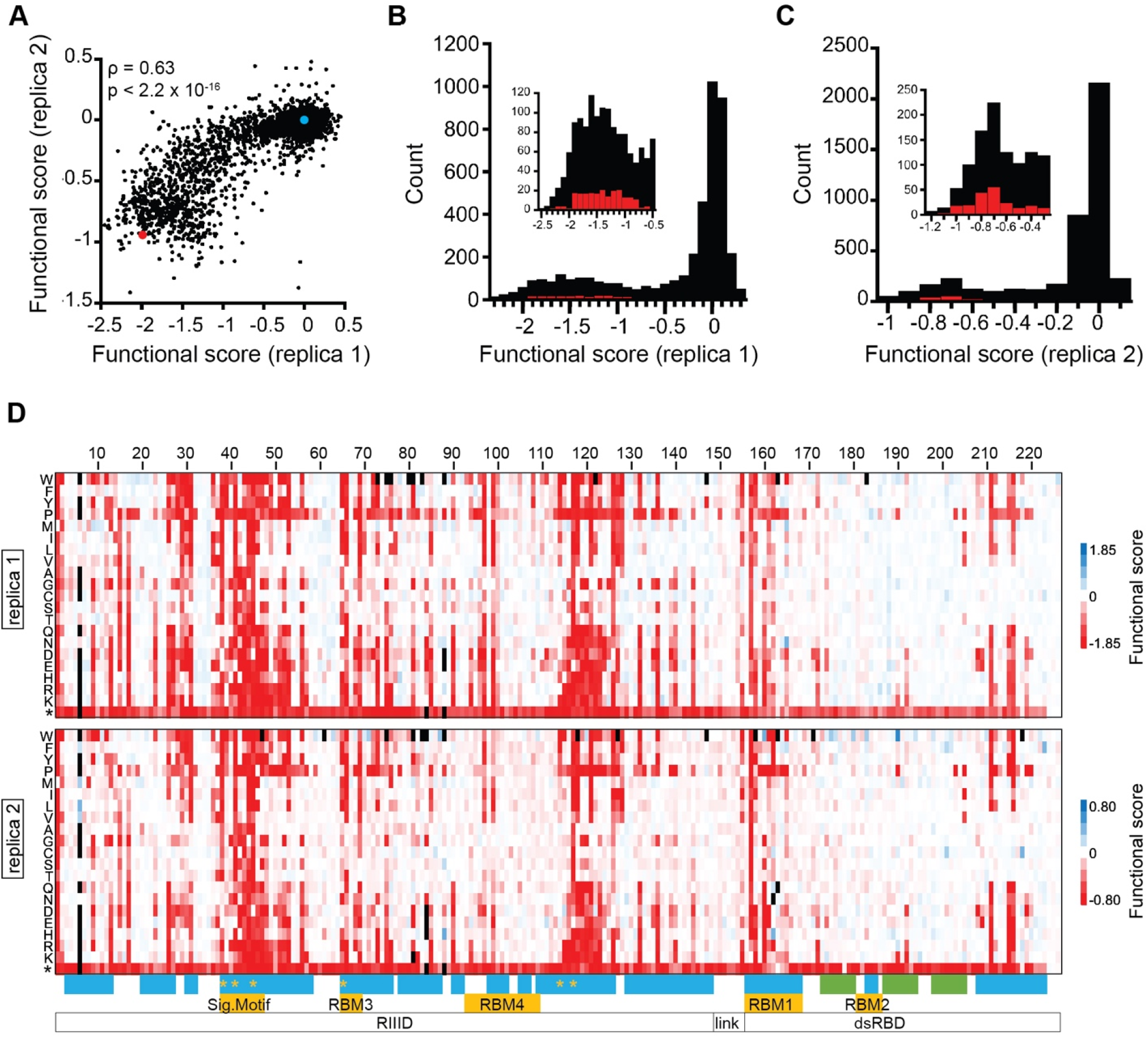
Correlation between functional scores of replica 1 and replica 2. (**A**) Despite differences in the range, functional scores between replica 1 and replica 2 correlate. The blue point represents wildtype, and the red point represents nonfunctional RNase III. Nonfunctional RNase III was determined by the weighted mean of all mutants with a non-functional start codon. (**B,C**) DFEs of functional scores for missense (black) and nonsense (red) mutations for each replica show a bimodal distribution with peaks at about 0 and -1.5 for replica 1 and 0 and -0.7 for replica 2. (**D**) Heat maps compare the functional scores from replica 1 and replica 2. Red is deleterious, white is neutral, blue is beneficial, and black indicates insufficient data due to low sequencing counts. The shading was set such that full red represented the average value corresponding to complete loss of RNase III (different scales in each replica, as indicated).

**Supplementary Figure S7:**
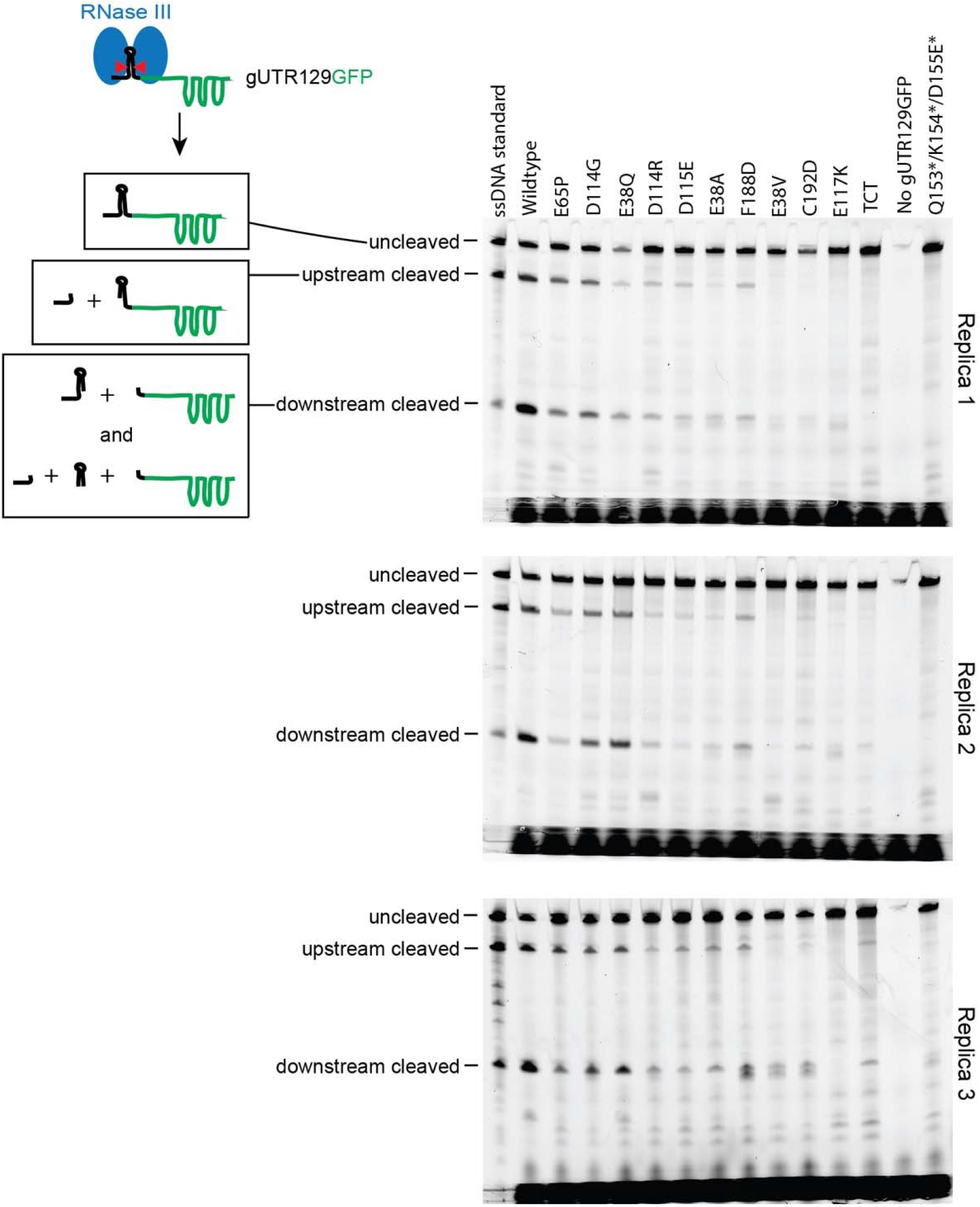
Representative primer extension gels for the mutants of Table 1. mRNA from cultures expressing each mutant in Table 1 was subjected to the fluorescent primer extension assay and analyzed by denaturing gel electrophoresis. gUTR129GFP cDNA products were compared to PCR-amplified ssDNA standards corresponding to the uncleaved (195 nt), upstream cleaved (155 nt), and downstream cleaved (73 nt) species.

**Supplementary Figure S8:**
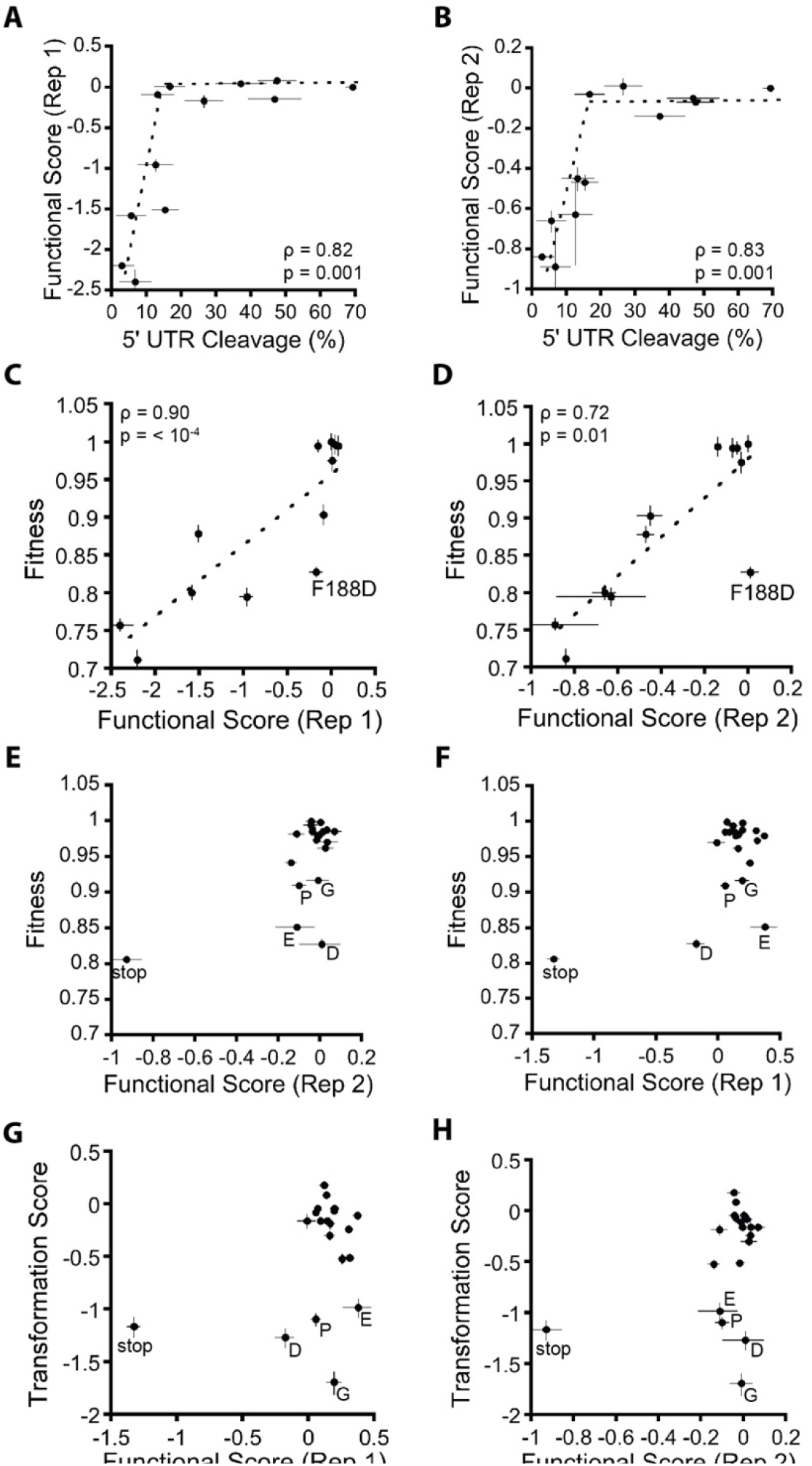
Analogous graphs of Figure 4 plotted using each replica of the functional score. The functional scores of replica 1 and replica 2 had different ranges. This figure provides graphs analogous to those in Figure 4 but using each functional score replica separately instead of using the weighted mean functional score. (**A,B**) Functional score as a function of percent 5’UTR cleavage. (**C,D**) Fitness as a function of functional score (**E,F**) Fitness as a function of functional score for mutations at F188 (**G,H**) Transformation score as a function of functional score for mutations at F188. Fitness was determined by measuring the growth rate relative to wildtype (error bars are the standard deviation *n*=3) except for panels E and F where the fitness values for all mutants except F188D are the weighted mean values from the DMS experiment. Primer extension assays were used to determine the % cleavage of the 5’UTR stem loop of *rnc* in gUTR129GFP (error bars are the standard deviation *n*=3). The fluorescence/OD of bacterial cultures reports on RNase III cleavage of the 5’UTR stem loop of *rnc* in gUTR129GFP (error bars are the standard deviation *n*=3). Dashed lines are linear fits except for panels A and B, in which case they are guides for the eye.

**Supplementary Figure S9:**
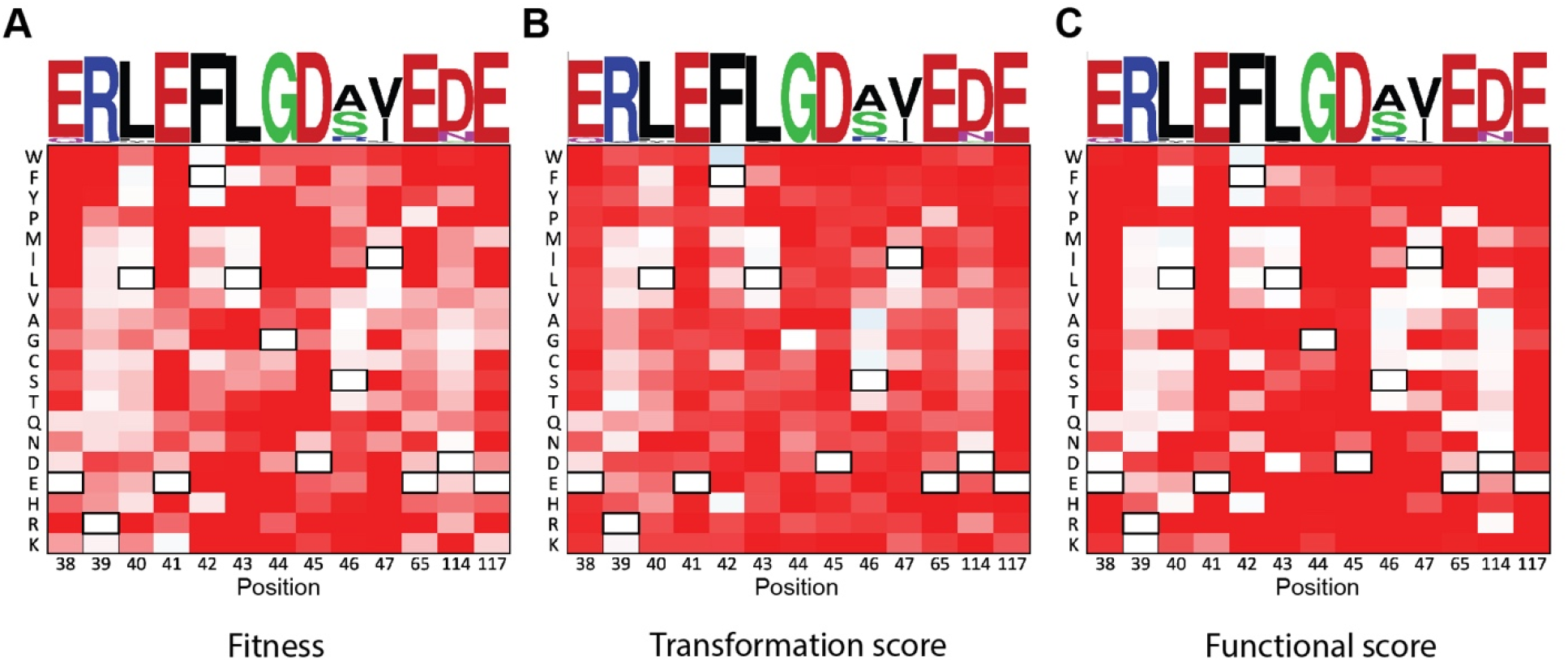
Fitness and functional effects of mutations in the RNase III signature motif and active site residues. The (**A**) weighted mean fitness, (**B**) transformation score, and (**C**) weighted mean functional score of significant residues within RNase III. Residues 38 – 47 are the RNase III signature motif that is highly conserved across all species. The sequence conservation of the RNase III family is indicated by the WebLogo at the top. Residues 38, 41, 45, 65, 114, and 117 are key active site residues. Black rectangles indicate the *E. coli* RNase III residue for each position.

## Captions for data files

**Data S1. Excel file for DMS sequencing counts, fitness values, and associated statistics for the fitness landscape**.

**Data S2. Excel file for DMS sequencing counts, transformation score values, and associated statistics for transformation scores**.

**Data S3. Excel file for DMS sequencing counts, functional score values, and associated statistics for functional scores**.

**Data S4. Excel file for *k\** values for sequence alignment, transformation score, and functional score**.

## Notes

### Competing Interest Statement

The authors have declared no competing interest.

